# A developmental transition in the sensory coding of limb kinematics in primary motor cortex

**DOI:** 10.1101/2020.12.14.422707

**Authors:** Ryan M. Glanz, James C. Dooley, Greta Sokoloff, Mark S. Blumberg

**Author notes:** Corresponding authors: Mark S. Blumberg, Ph.D., Ryan M. Glanz. Lead Contact: Mark S. Blumberg.

## Abstract

Primary motor cortex (M1) undergoes protracted development in mammals, functioning initially as a sensory structure. Throughout the first postnatal week in rats, M1 is strongly activated by self-generated forelimb movements—especially by the twitches that occur during active sleep. Here, we quantify the kinematic features of forelimb movements to reveal receptive-field properties of individual units within the forelimb region of M1. At postnatal day (P) 8, nearly all units were strongly modulated by movement amplitude, especially during active sleep. By P12, only a minority of units continued to exhibit amplitude-tuning, regardless of behavioral state. At both ages, movement direction also modulated M1 activity, though to a lesser extent. Finally, at P12, M1 population-level activity became more sparse and decorrelated, along with a substantial alteration in the statistical distribution of M1 responses to limb movements. These findings reveal a transition toward a more complex and informationally rich representation of movement long before M1 develops its motor functionality.

## Introduction

Despite its name, primary motor cortex (M1) is increasingly appreciated as an integrator of sensory input [1,2]. The sensory-processing capabilities of M1 are especially evident in early development before it plays any role in the production of movement. At such ages, M1 functions exclusively as a prototypical sensory structure [3-5], with M1 activity increasing after—not before—movements have been generated, indicative of reafference [3,4,6]. M1’s motor capabilities, on the other hand, emerge gradually: Electrical stimulation of M1 does not reliably produce movement until postnatal days (P) 25–30 [7], and the precise age at which M1 activity begins to precede self-generated movements remains unknown. Before P25, self-generated movements are produced exclusively by brainstem structures, including the red nucleus [8,9]. Based in part on the somatotopic alignment between M1’s early sensory map and its later motor map, it seems clear that the former provides a structural foundation for the latter [4]. This structural foundation may subsequently serve as a “ground truth” reference by which emerging motor outflow is directed to its correct efferent target.

In early development, reafferent activation of M1 neurons is modulated by behavioral state. Reafference arising from myoclonic twitches—the jerky movements of the limbs and whiskers that occur abundantly during active sleep—robustly and reliably triggers M1 activity, whereas sustained bouts of wake movements result in only weak activation [3,4,6]. Because myoclonic twitches are brief, discrete events that occur against a background of muscle atonia, they are well-positioned to convey high-fidelity proprioceptive information to M1. By comparison, limb movements during wake are more sustained than twitches and typically involve the simultaneous engagement of multiple muscle groups within and across limbs. As demonstrated with the forelimbs in P8 rats, reafferent activity during wake is inhibited early in the processing stream at the level of the external cuneate nucleus [4,6], perhaps to prevent a saturated reafferent signal from muddying downstream sensory representations that are still developing. Given these unique features of twitching and the fact that cortical development is an activity-dependent process [10-12], we have proposed that twitches help to establish and refine M1’s somatotopically precise sensory map [4].

In adult monkeys, activity in M1 is correlated with the amplitude of the movements that it produces [13]. In infant rats, activity in primary somatosensory cortex (S1) is similarly correlated with the amplitude of whisker movements: Specifically, we recently reported in P5 rats that high-amplitude whisker movements reliably trigger greater activation of S1 barrel cortex than low-amplitude whisker movements [14]. Building on the notion that M1 initially functions as a canonical sensory structure, we predicted that it should also exhibit evidence of amplitude-dependent sensory coding. To test this prediction, we utilized recent advances in machine vision tools (DeepLabCut; [15,16]) to track the trajectory of limb movements while simultaneously recording extracellular activity in the forelimb region of M1. As predicted, we demonstrate that M1 activity at P8 is robustly tuned to the amplitude of forelimb movements during active sleep, but not wake. Moreover, nearly all M1 units share similar amplitude-tuning curves, leading to a repetitive—and thus, redundant— coding of movement amplitude at P8. In contrast, at P12, only a minority of M1 units continue to exhibit amplitude-tuning, regardless of sleep-wake state. M1 units also exhibit sensitivity to movement direction at both ages, although this feature of M1 tuning is less robust than that of amplitude. “Background” neural activity (occurring during periods of behavioral quiescence) increases dramatically over this period, marking a key transition from discontinuous to continuous activity. This transition is also characterized by increased sparsity and entropy across units, suggesting that M1 activity at P12 carries more informational content. Indeed, redundant “all-or-nothing” responses across M1 units, triggered by forelimb movements, are common at P8, and become increasingly rare at P12, further increasing the informational content embedded within M1 activity. All together, these results highlight multiple transitional features in M1’s developmental cascade from an exclusively sensory structure to one that will ultimately integrate sensorimotor information to organize motor outflow and support motor learning [17,18].

## Results

To characterize the relationship between movement kinematics and M1 activity, we tracked forelimb movements in three dimensions for one hour using two high-speed (100 frames/s) cameras oriented perpendicularly to each other (Fig 1A). At the same time, we performed extracellular recordings in the contralateral forelimb region of M1 (Fig 1B–C). Because preliminary analyses indicated similar results for single- and multi-units, all units were analyzed together. Fig 1D depicts representative behavioral and electrophysiological data collected at each age; most striking is the transition from discontinuous to continuous unit activity from P8 to P12.

**Fig 1.**
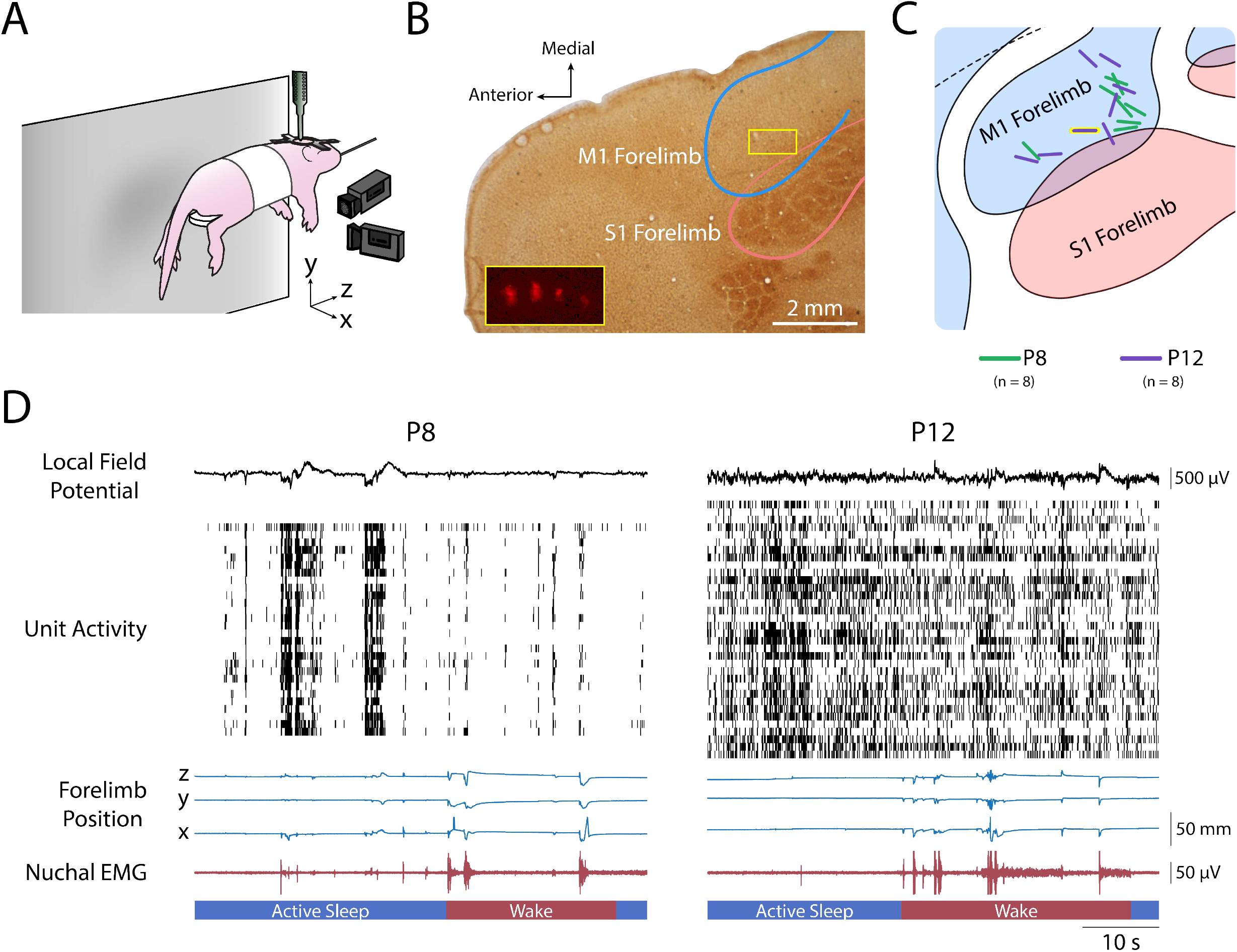
Experimental setup, histology, and representative data. (**A**) P8 and P12 rats were head-fixed with their torso supported and their limbs free to dangle. Two cameras were placed perpendicularly: One front-view camera captured the x-dimension (medial-lateral) of right forelimb movements, and one side-view camera captured the y-dimension (dorsal-ventral) and z-dimension (anterior-posterior). A 4×4 silicon depth electrode was lowered into the forelimb region of contralateral M1 to record extracellular activity. (**B**) Photomicrograph of electrode placement in M1. Cortical tissue was flattened before sectioning, followed by staining with cytochrome-oxidase to reveal the primary somatosensory cortical representations. The M1 and S1 forelimb boundaries are depicted as blue and pink lines, respectively. The yellow box in the M1 forelimb region delineates the location of the four-shank electrode for this pup. Inset: magnified view of the four-shank electrode revealed using fluorescence. (**C**) Electrode placements for the eight subjects at P8 and P12 represented as green and purple lines, respectively. The location of the electrode in (B) is shown with a yellow border. (**D**) Representative data from a P8 (left) and P12 (right) rat. From top to bottom: the local field potential (LFP) in M1; single- and multi-unit activity in M1, with each row denoting a different single-unit or multi-unit and each vertical tick denoting an action potential; traces of right forelimb position in the x-, y-, and z-dimensions; nuchal EMG activity; and active sleep (blue bars) and wake (red bars).

The trajectory of forelimb movements was quantified using DeepLabCut (DLC), a machine-vision tool for behavioral analysis [15,16]. Movement amplitude of the forelimb was aggregated across three dimensions before analysis (Fig 2A). The amplitude of forelimb twitches was exponentially distributed at P8 and P12, with small twitches being more frequent than large twitches (Fig 2B). In contrast, the amplitude of wake movements followed a positively skewed normal distribution. Whereas small forelimb movements resulted primarily in displacement of the distal forelimb (i.e., wrist and digits; S1A Fig), large forelimb movements resulted in greater displacement of the proximal forelimb (i.e., forearm and elbow). Finally, movement velocity and acceleration were highly correlated with movement amplitude and thus were not analyzed further (S1B Fig).

**Fig 2.**
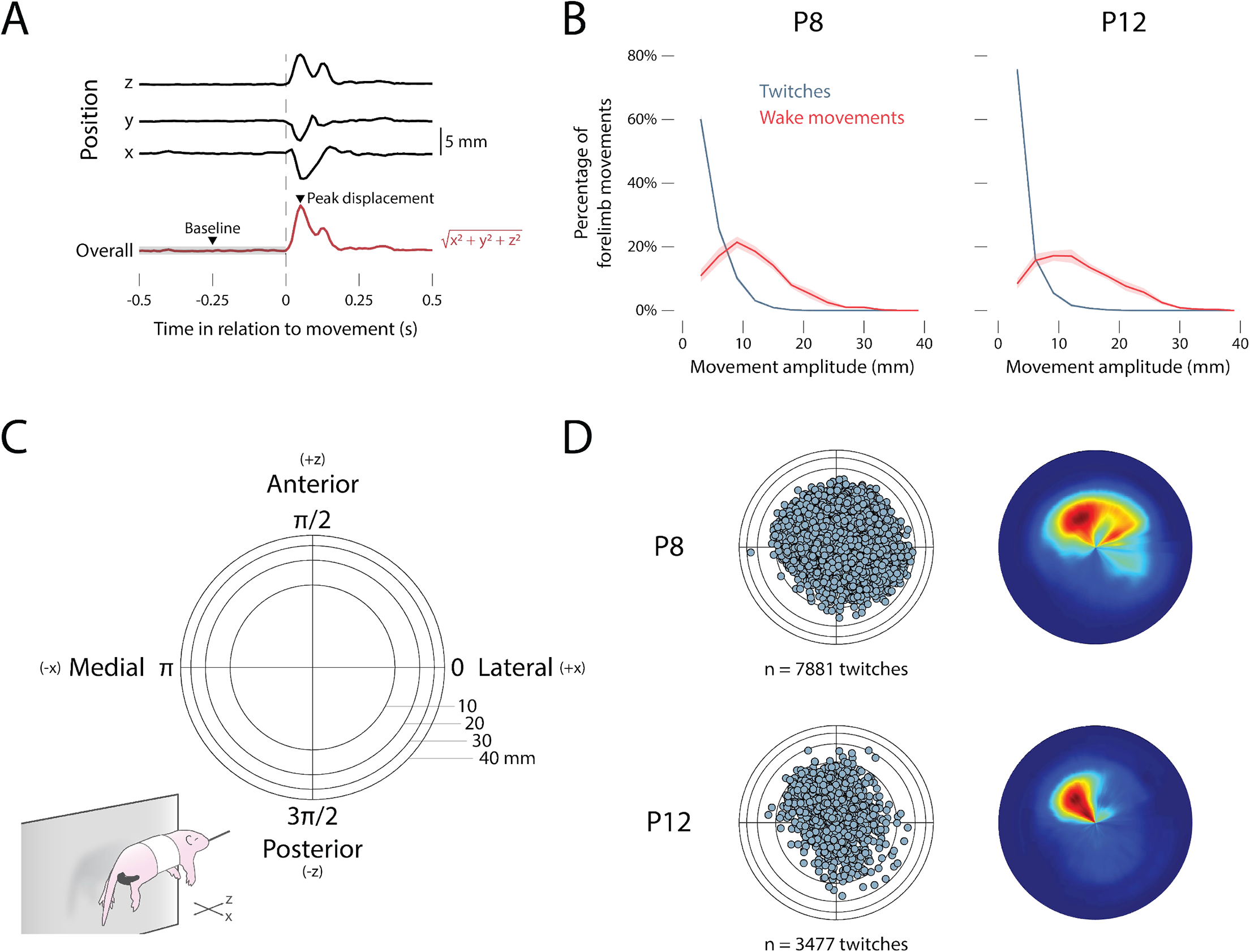
Kinematics of forelimb movements across behavioral state. (**A**) Depiction of how forelimb movement amplitude was calculated. For each detected twitch and wake movement, the position across the x-, y-, and z-dimensions was summed using the Pythagorean theorem. Movement amplitude was defined as the difference between the peak position of the forelimb 0–250 ms after a movement and the median position of the forelimb at baseline (i.e., -0.5 to 0 s before a movement). (**B**) Mean (±SEM) of the percentage of forelimb twitches (blue) and wake movements (red) as a function of amplitude at P8 (left) and P12 (right). (**C**) Two-dimensional representation of twitch direction in polar space. The x-dimension (medial-lateral) and the z-dimension (anterior-posterior) are shown. (**D**) Left: Scatterplot showing the position of the right forelimb at peak twitch displacement for all twitches at P8 (top) and P12 (bottom). Right: Heatmap showing the most common position of the right forelimb at peak twitch displacement for all twitches at P8 and P12. See also S1 and S2 Figs.

Because the pup’s limbs dangled freely in our testing environment, forelimb twitches traveled in a pendular motion that was skewed heavily in the positive y-dimension (see S1C–D Fig). As a consequence, displacement in the y-dimension did not provide unique information about movement direction. Therefore, we confined our directional analyses to movements in the x- and z-dimensions (anterior-posterior and medial-lateral, respectively; Fig 2C).

Forelimb twitches occurred in all directions relative to the limb’s resting position (Fig 2D) but were most likely to travel anteriorly at P8 and anteriorly and medially at P12. Movement direction could not be assessed accurately for wake movements because such movements are typically produced in rapid succession with many direction changes.

### M1 activity is strongly modulated by behavioral state

Over the 60-min recording periods, P8 and P12 rats spent 57.4 ± 3.2% and 44.8 ± 7.6% of the time in active sleep, respectively (t(14) = 1.51, p = .153, adj. 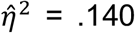; S2A Fig). Twitches occurred significantly more frequently than wake movements at both ages (P8: F(1, 7) = 92.08, p < .001, adj. 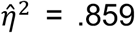; P12: F(1, 7) = 7.11, p = .018, adj. 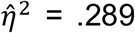; S2B Fig). The relative abundance of active sleep and twitches, especially at P8, provides ample opportunity for reafference from twitching limbs to drive M1 activity.

Consistent with previous reports [4,6], we found here at P8 that twitches drove relatively more M1 activity than did wake movements (S2C Fig). This state-dependence in reafferent activity was significant across age (F(1, 14) = 30.16, p < .001, adj. 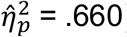). Compared with previous studies that relied on EMG to detect movement [3,4,6], the use here of a video-based method substantially increased the number of twitches and wake movements that were detected. As a consequence, at P8 we now reliably detected more M1 activation in response to wake movements than previously reported [4,6]; nonetheless, reafferent responses to twitches were still larger than they were to wake movements. Moreover, at both ages, twitches were significantly more likely than wake movements to trigger M1 activity (P8: F(1, 7) = 21.25, p = .002, adj. 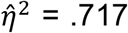; P12: F(1, 7) = 17.21, p = .004, adj. 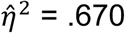; S2D Fig).

### M1 units are tuned to movement amplitude at P8

As mentioned above, we recently reported in P5 rats that whisker-movement amplitude reliably predicts activity in S1 barrel cortex [14]. Accordingly, we hypothesized here that M1 similarly codes for movement amplitude during the sensory stage of its development. To test this hypothesis, twitches and wake movements were sorted into amplitude bins from 0–16 mm. Amplitude bins were scaled logarithmically in increments from 2^0^ to 2^4^ mm to account for the exponential distribution of twitches. To avoid potential bias, equal numbers of anterior/posterior and medial/lateral movements were selected at random for these analyses.

At P8, twitches of increasing amplitude triggered increasing unit activation in M1 (Fig 3A). This amplitude-tuning was not observed for wake movements at P8, nor for twitches or wake movements at P12. Fig 3B quantifies the relationship between movement amplitude and *response strength*—the difference between the mean firing rate 0–250 ms after a movement and the mean baseline firing rate determined at -3 to -2 s before a movement. At P8, response strength significantly increased with respect to increasing twitch amplitude, but not wake-movement amplitude (F(1, 216) = 250.27, p < .001, adj. 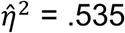). This was not the case at P12 (F(1, 249) = 0.42, p = .520, adj. 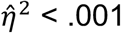).

**Fig 3.**
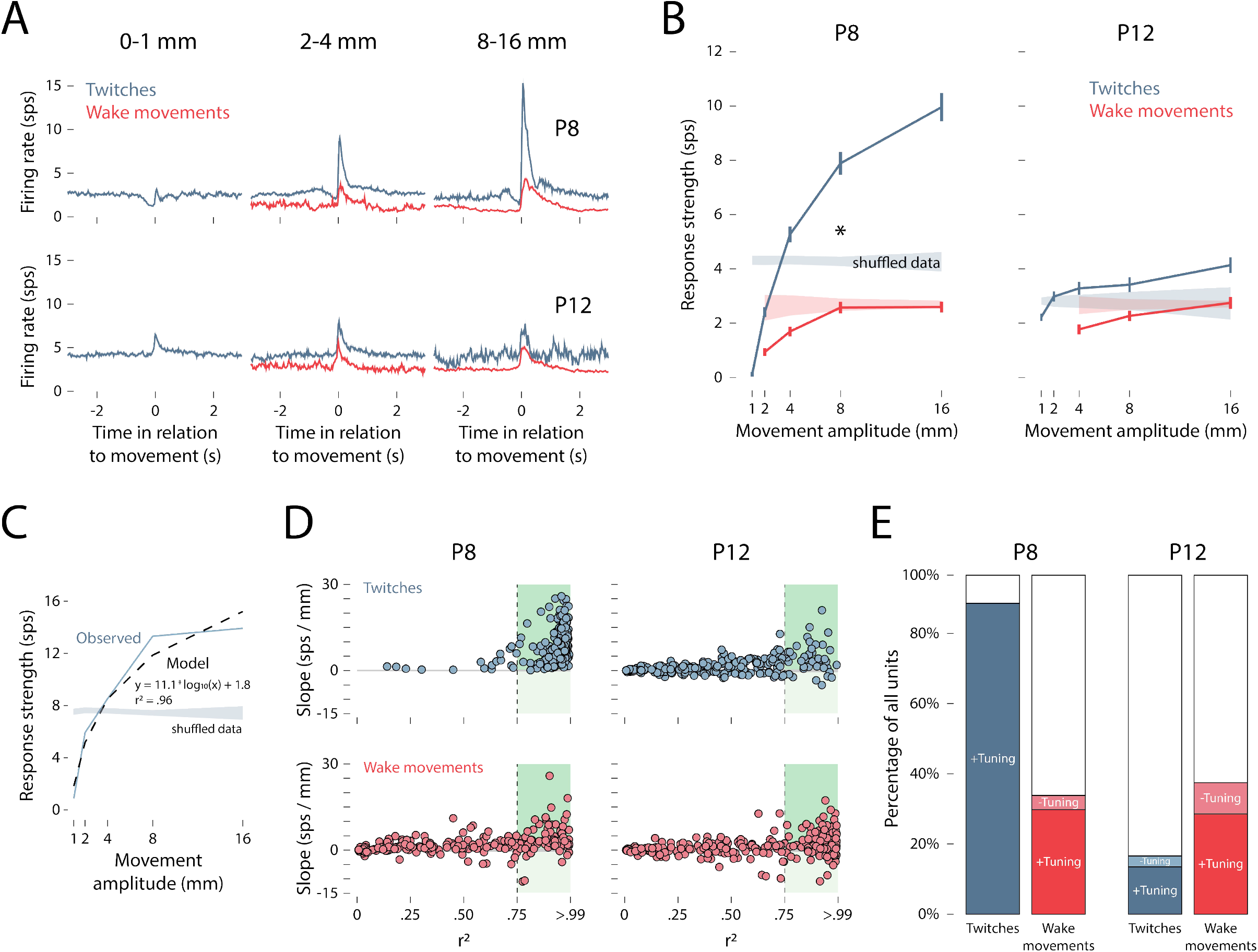
Relationship between movement amplitude and M1 unit activity. (**A**) Peristimulus time histograms of mean firing rate of M1 units in relation to the onset of twitches (blue) and wake movements (red) at P8 (top row) and P12 (bottom row). From left to right, the columns show small-amplitude (0–1 mm), medium-amplitude (2–4 mm), and large-amplitude (8–16 mm) forelimb movements. (**B**) Mean (±SEM) response strength (Δfiring rate in relation to baseline) for twitches (blue) and wake movements (red) for all M1 units at P8 (left) and P12 (right). The right-edge of each amplitude bin is labeled on the x-axis. Color-coded shaded regions indicate 99% confidence intervals based on shuffled data. Asterisk indicates significant difference between twitches and wake movements at P8 (p < .05). (**C**) Representative example of an individual M1 unit’s relationship between response strength and movement amplitude at P8, and its fit to a logarithmic model. The observed data (blue solid line) and model data (black dashed line) are shown alongside the 99% confidence interval based on shuffled data (blue shaded region). The slope of the model (11.1 sps/mm) indicates the strength of the unit’s relationship with movement amplitude, and the r2 value (.96) represents the goodness-of-fit to the logarithmic function. (**D**) Applying the method in (C) to all M1 units, the slope and r2 value for each unit is shown for twitches (blue) and wake movements (red) at P8 and P12. The gray-shaded horizontal lines represent the 99% confidence intervals based on shuffled data, and the dashed vertical lines represent an r2 threshold of .75. The shaded green regions to the right of the .75 threshold show units that are positively (slope > 0) and negatively (slope < 0) correlated with movement amplitude. (**E**) Based on the r2 threshold of .75 used in (D), stacked plots show the percentage of units at P8 and P12 that are positively tuned to movement amplitude for twitches (dark blue) and wake movements (dark red), negatively tuned for twitches (light blue) and wake movements (light red), or not tuned (white). See also S3 Fig.

Although wake movements at P8, and all movements at P12, did not show a significant relationship with amplitude when averaged across units, it is still possible that some individual units were tuned to movement amplitude. To characterize the relationship between movement amplitude and response strength on a unit-by-unit basis, each unit’s response strength was fit to a logarithmic model with respect to movement amplitude (Fig 3C). A logarithmic model was chosen based on the relationship between twitch amplitude and response strength observed at P8. Both the slope and the goodness-of-fit (r^2^) of the model were used to assess each unit’s relationship with movement amplitude. In Fig 3D, the r^2^ value and slope of the model for each M1 unit are shown. A minimum r^2^ value of .75 was chosen to consider an individual unit to be tuned to movement amplitude. At P8, the vast majority of M1 units showed positive tuning to amplitude, with r^2^ values greater than .75 and slopes that exceeded the 99% confidence interval of shuffled data. In contrast, for wake movements at P8 and all movements at P12, only a fraction of M1 units met these criteria, and some units even displayed negative relationships with amplitude (i.e., negative tuning).

As shown in the stacked plots in Fig 3E, nearly all M1 units at P8 exhibited a positive tuning to twitch amplitude (positive: 91.7%; negative: 0%); far fewer units exhibited amplitude-tuning to wake movements (positive: 30.9%; negative: 4.2%). At P12, relatively few units exhibited tuning to twitch amplitude (positive: 14.0%; negative: 3.2%) or wake-movement amplitude (positive: 29.6%; negative: 9.2%). Finally, to confirm that these findings were not driven by an arbitrary selection of an r^2^ threshold of .75, we repeated these tests using thresholds of .50 and .90 and found the same pattern of results (S3A Fig).

When positively and negatively tuned units are considered separately, only twitches showed an age-related change in tuning strength: Small twitches (0–2 mm) at P12 were better able than small twitches at P8 to drive M1 activity (F(1.31, 291.02) = 10.01, p = .001, adj. 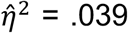; S3B Fig). In contrast, amplitude-tuning for wake movements did not change with age for either positively or negatively tuned units. When response strength was standardized (S3C Fig), M1 units at P8 showed highly redundant responses to increasing twitch amplitude, but not wake-movement amplitude. At P12, M1 units did not show redundant responses to movement amplitude, regardless of sleep-wake state.

In summary, these findings extend previous reports [4,6] by showing that forelimb twitches in early development differentially trigger M1 activity across a range of movement amplitudes. This is the first demonstration of amplitude coding in infant M1 and is consistent with similar findings in S1 barrel cortex [14].

### M1 units are sensitive to movement direction

We next analyzed M1 activity as a function of twitch direction. (Because wake movements occur in prolonged bouts with multiple changes in direction, they could not be included in this analysis.) Twitch direction was analyzed separately along two dimensions: Anterior-posterior and medial-lateral (Fig 4A). Again, to avoid potential bias, we ensured that twitches in each direction had identical amplitude distributions. Twitches of the forelimb in each direction produced similar responses in M1 units at both P8 and P12 (Fig 4B). However, as shown in Fig 4C, the average response strength at P8 was significantly lower for anterior compared to posterior twitches (F(1, 216) = 51.05, p < .001, adj. 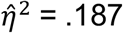) and medial compared to lateral twitches (F(1, 216) = 53.75, p < .001, adj. 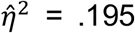). At P12, there was a small but statistically significant difference in response strength between anterior and posterior twitches (F(1, 249) = 9.80, p = .002, adj. 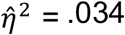), but not between medial and lateral twitches (F(1, 249) = 0.28, p = .599, adj. 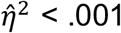).

**Fig 4.**
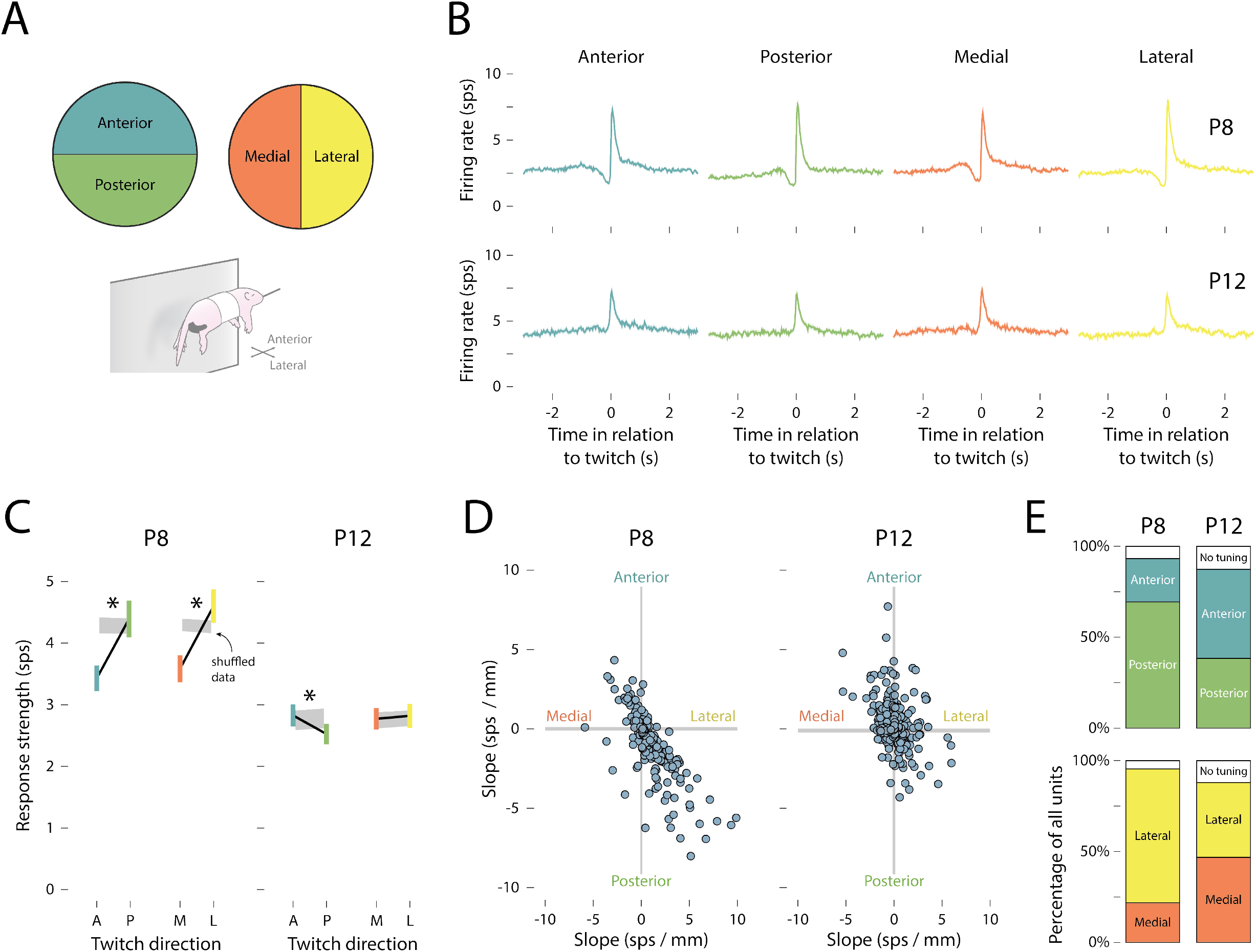
Effect of movement direction on M1 unit activity. (**A**) Individual forelimb twitches were classified as moving in anterior (blue), posterior (green), medial (orange), or lateral (yellow) directions. (**B**) Peristimulus time histograms of the mean (±SEM) firing rate of M1 units in relation to the onset time of (from left to right) anterior, posterior, medial, and lateral forelimb twitches at P8 (top) and P12 (bottom). (**C**) Mean (±SEM) response strength (Δfiring rate in relation to baseline) for anterior (blue) and posterior (green) twitches, and for medial (orange) and lateral (yellow) twitches at P8 (left) and P12 (right). The gray shaded regions indicate 99% confidence intervals based on shuffled data. Asterisks indicate a significant difference between movement directions (p < .05). (**D**) The slope (strength of the unit’s relationship with movement direction) for individual units is represented along the medial-lateral (x) and anterior-posterior (y) axes. The gray shaded horizontal and vertical lines indicate 99% confidence intervals based on shuffled data. (**E**) Top row: Stacked plots show the percentage of units tuned to anterior (blue) or posterior (green) twitches at P8 and P12. Bottom row: Stacked plots show the percentage of units tuned to medial (orange) or lateral (yellow) twitches at P8 and P12. The white regions indicate untuned units.

Again, the trends in direction-related response strength, averaged across all units, may have obscured the direction-tuning of individual units. Accordingly, the direction-tuning of individual units was assessed by measuring the difference in response strength between anterior/posterior and medial/lateral twitches. At P8, individual units tended to be responsive to twitches that traveled posteriorly and laterally (Fig 4D). By P12, receptive fields were more uniformly distributed across all four direction combinations. Indeed, at P8, the majority of units were tuned to posterior rather than anterior movements (69.6% and 24.0%, respectively) and to lateral rather than medial movements (73.3% and 22.1%, respectively; Fig 4E). At P12, however, direction-tuning was more evenly distributed (range: 38.4–49.2%). Thus, as with amplitude tuning, there was an age-related reduction among M1 units in the redundancy of direction tuning.

### Sparse background activity emerges at P12

Thus far, we have focused on the correlation between movement kinematics and the firing rate of individual units. However, the activity *across* units—that is, population spiking activity—also undergoes a marked developmental shift: Population activity shifts from discontinuous at P8 to continuous at P12 (Fig 5A). To characterize this change in population activity, we draw a distinction between <u>movement-related activity</u>—M1 activity occurring during and immediately after (i.e., within 250 ms) limb movements—and <u>background activity</u>—M1 activity occurring during periods in which limb movements are absent.

**Fig 5.**
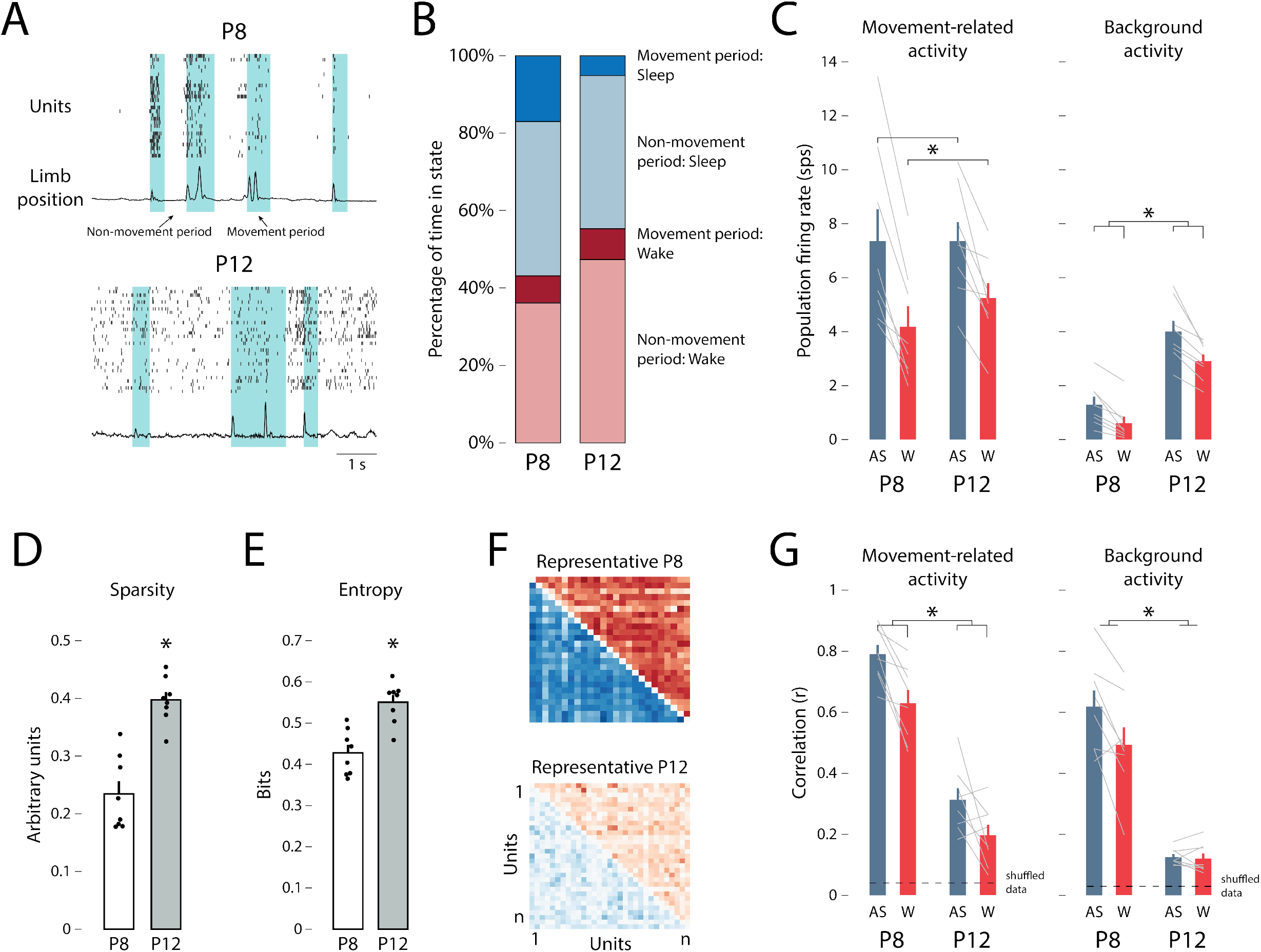
Population spiking activity decorrelates between P8 and P12. (**A**) Population spiking activity and limb position from a representative P8 (top) and P12 (bottom) rat are shown. A spike raster (each row represents one unit) and limb position trace (normalized) is shown for each pup. Green-shaded regions represent movement periods, and the unshaded regions represent non-movement periods. (**B**) The percentage of time spent in movement or non-movement periods (Movement period (sleep): blue; Non-movement period (sleep): light blue; Movement period (wake): red; Non-movement period (wake): light red) is shown for P8 (left) and P12 (right) animals. (**C**) Mean (±SEM) movement-related activity (left-hand plot) and background activity (right-hand plot) firing rate for active sleep (blue) and wake (red) for P8 (left) and P12 (right). Firing rates for each rat are shown as gray lines. Asterisks denote a significant main effect of behavioral state (left-hand plot; p < .05) and significant main effects of behavioral state and age (right-hand plot; p < .05). (**D**) Mean (±SEM) values of sparsity for P8 (white) and P12 (gray) rats. Black dots show the values for individual pups. Asterisk indicates significant difference between P8 and P12 (p < .05). (**E**) Mean (±SEM) values of entropy for P8 (white) and P12 (gray) rats. Black dots show the values for individual pups. Asterisks indicates significant difference between P8 and P12 (p < .05). (**F**) Representative correlation matrices for each unit-unit pair (x- and y-axes) for a P8 (top) and P12 (bottom) rat during movement periods. Blue squares indicate correlations during active sleep and red squares indicate correlations during wake. Darker colors denote higher r2 values. (**G**) Mean (±SEM) correlation coefficient (r) during active sleep (blue) and wake (red) for P8 (left) and P12 (right) animals for movement-related activity (left-hand plot) and background activity (right-hand plot). Correlation coefficients for each rat are shown as gray lines. Dashed horizontal lines indicate 99% confidence interval based on shuffled data. Asterisks represent significant main effects of behavioral state and age (left-hand plot; p < .05) and a significant main effect of age (right-hand plot; p < .05).

To characterize the change in population spiking activity between P8 and P12, we first characterized four periods during each recording session: (1) movement periods during active sleep (i.e., periods of twitches), (2) non-movement periods during active sleep, (3) movement periods during active wake (i.e., periods of wake movements) and, (4) non-movement periods during wake (Fig 5B). Pups at both ages spent a greater percentage of time in non-movement periods (P8: AS = 39.9%, W = 36.1%; P12: AS = 39.5%, W = 47.4%) than in movement periods (P8: AS = 17.0%, W = 7.0%; P12: AS = 5.2%, W = 8.0%). As shown in Fig 5C, movement-related population activity was significantly higher during active sleep than wake (F(1, 14) = 53.97, p < .001, adj. 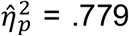), and this state difference did not change between P8 and P12 (F(1, 14) = 0.22, p = .644, adj. 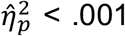). Conversely, background population activity was not only higher during active sleep compared with wake (F(1, 14) = 47.31, p < .001, adj. 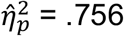), but exhibited a three- to four-fold increase between P8 and P12 (F(1, 14) = 39.21, p < .001, adj. 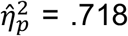). Thus, the increase in M1’s population spiking activity between P8 and P12 occurred almost entirely during non-movement periods.

Next, we asked whether the patterning of M1 population activity changed between P8 and P12. We first measured the change in sparsity (the degree to which action potentials are uniformly distributed across time) and entropy (the informational capacity available given the patterning of activity) from P8 to P12. Importantly, sparsity and entropy interact to efficiently encode sensory features in cortical networks [19]. Indeed, across M1 units, both sparsity and entropy increased significantly between P8 and P12 (sparsity: t(10.89) = 5.97, p < .001, adj. 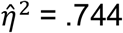; entropy: t(14) = 4.71, p < .001, adj. 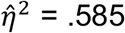; Fig 5D–E).

To assess whether the increase in uniform background activity affected population dynamics in M1, we computed unit-by-unit correlations of activity during each of the four periods outlined above. Fig 5F shows correlation matrices for representative pups at P8 and P12 during active sleep (blue) and wake (red) movement periods. Movement-related activity, but not background activity, was significantly more correlated during active sleep than wake (F(1, 14) = 28.02, p < .001, adj. 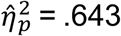; Fig 5G). Moreover, movement-related activity was significantly less correlated at P12 than at P8 (F(1, 14) = 91.97, p < .001, adj. 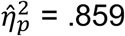; Fig 5G), as was background activity (F(1, 14) = 53.41, p < .001, adj. 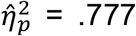). This decorrelation of population activity further indicates a transition toward sparse—and therefore less redundant—M1 activity at P12. Although population activity was significantly less correlated at P12, the observed correlations were still stronger than expected (i.e., compared with shuffled data). In other words, population spiking activity at P12 continued to exhibit an organized temporal structure.

Next, we sought to characterize the population-level activity of M1 units in response to forelimb movements. For each given movement, an M1 unit was considered “responsive” if its firing rate increased significantly (i.e., exceeded the 95% confidence interval) relative to its baseline firing rate. If this threshold was not reached, the M1 unit was considered “unresponsive” to that particular movement. For each limb movement, then, the percentage of responsive M1 units— the *population response*—was calculated (Fig 6A). Fig 6B plots the mean percentage of forelimb movements across pups (y-axis) that triggered a given population response (x-axis).

**Fig 6.**
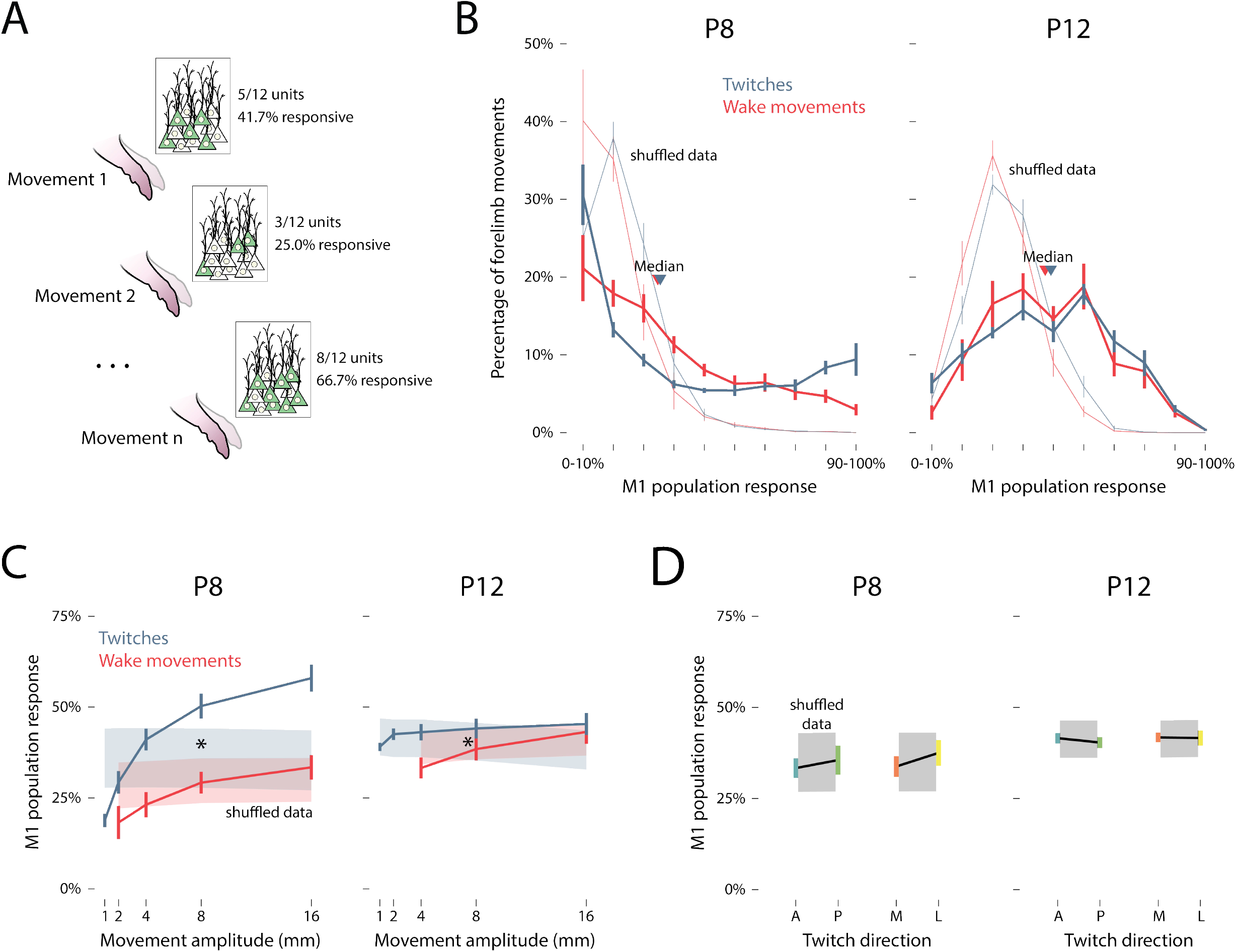
Population responses in M1. (**A**) Illustration of the relationship between forelimb movements (1 through n) and percentage of M1 units responding to a given movement (i.e., population response). (**B**) Mean (±SEM) percentage of twitches (blue) and wake movements (red) that yielded population responses of 0–10% to 90–100% at P8 (left) and P12 (right). Median values for twitches (blue arrows) and wake movements (red arrows) are also shown. Light blue and red lines indicate shuffled data for twitches and wake movements, respectively. (**C**) Mean (±SEM) population response for M1 units as a function of the amplitude of twitches (blue) and wake movements (red) at P8 (left) and P12 (right). Color-coded shaded regions represent 99% confidence intervals based on shuffled data. Asterisks indicates significant main effect of twitches versus wake movements (p < .05). (**D**) Mean (±SEM) population response of M1 units for anterior (blue), posterior (green), medial (orange), and lateral (yellow) twitches at P8 (left) and P12 (right). Gray-shaded regions indicate 99% confidence intervals based on shuffled data.

At P8, a plurality (20-30%) of forelimb movements during sleep and wake triggered only a small percentage of M1 units (Fig 6B, left). Also, twitches, but not wake movements, regularly triggered activity (∼10% of responses) in nearly all M1 units. As a consequence, population responses at P8 followed a near-exponential distribution, with “all-or-nothing” responses occurring frequently—especially in response to forelimb twitches.

At P12, the distributions of the population responses to twitches and wake movements changed substantially (Fig 6B, right). Overall, forelimb movements at this age tended to trigger activity in approximately half of the M1 units, resulting in a roughly normal distribution. Indeed, all eight of the P12 rats—but only two of the eight P8 rats—exhibited normal distributions of their population responses (see S4 table). Population responses were significantly larger for twitches than wake movements, but this effect was smaller than that of age (P8 vs. P12: F(3.51, 49.15) = 26.93, p < .001, adj. 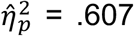; AS vs. W: F(1, 14) = 4.63, p = .049, adj. 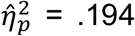). Furthermore, the distributions of population responses shifted as “all-or-nothing” responses (i.e., <10% or >90% of units responding) decreased from 26.9% at P8 to just 1.4% at P12. In other words, M1 population responses became substantially less redundant by P12.

We next determined whether movement kinematics predict M1 population responses. As shown in Fig 6C, the population response increased significantly as a function of movement amplitude at both ages, but the state-dependence of this relationship was stronger at P8 (F(1, 7) = 22.06, p < .001, adj. 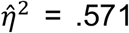) than at P12 (F(1, 7) = 9.37, p = .008, adj. 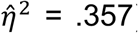). Overall, larger movements drove more activity within and across M1 units, especially during active sleep at P8.

In contrast with movement amplitude, twitch direction was not significantly related to population response at either age (anterior vs. posterior: F(1, 14) = 2.26, p = .155, adj. 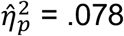; medial vs. lateral: F(1, 14) = 2.70, p = .123, adj. 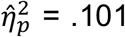; Fig 6D). This finding mirrors the small (though significant) differences in direction-tuning observed in individual M1 units at P8.

In summary, we found that M1 background activity increased sharply between P8 and P12, and that all population activity became more sparse and decorrelated across these ages. Also, population responses became less sensitive to movement amplitude at P12, and at the same time displayed a substantially different statistical distribution (i.e., normal versus exponential) in response to self-generated movements.

### Kinematic tuning is not mediated by spindle bursts

Spindle bursts are brief thalamocortical oscillations (10–20 Hz; Fig 7A–B) that are thought to contribute to early cortical development [14,20-22]. Because spindle bursts are readily detectable at P8 but not P12 [23], they could potentially mediate the effect of twitch amplitude on M1 activity at that earlier age. Indeed, spindle bursts were more likely to occur during active sleep than during wake (t(7) = 2.94, p = .022, adj. 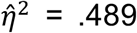; Fig 7C). However, whereas twitches and wake movements occurred at a rate of 28.3 ± 4.6 min^-1^ and 11.7 ± 1.2 min^-1^, respectively, spindle bursts occurred at 4.0 ± 0.7 min^-1^ during active sleep and 2.5 ± 0.4 min^-1^ during wake. Consequently, only a small percentage of forelimb movements (10.0 ± 1.4%) occurred within ±0.5 s of a spindle burst.

**Fig 7.**
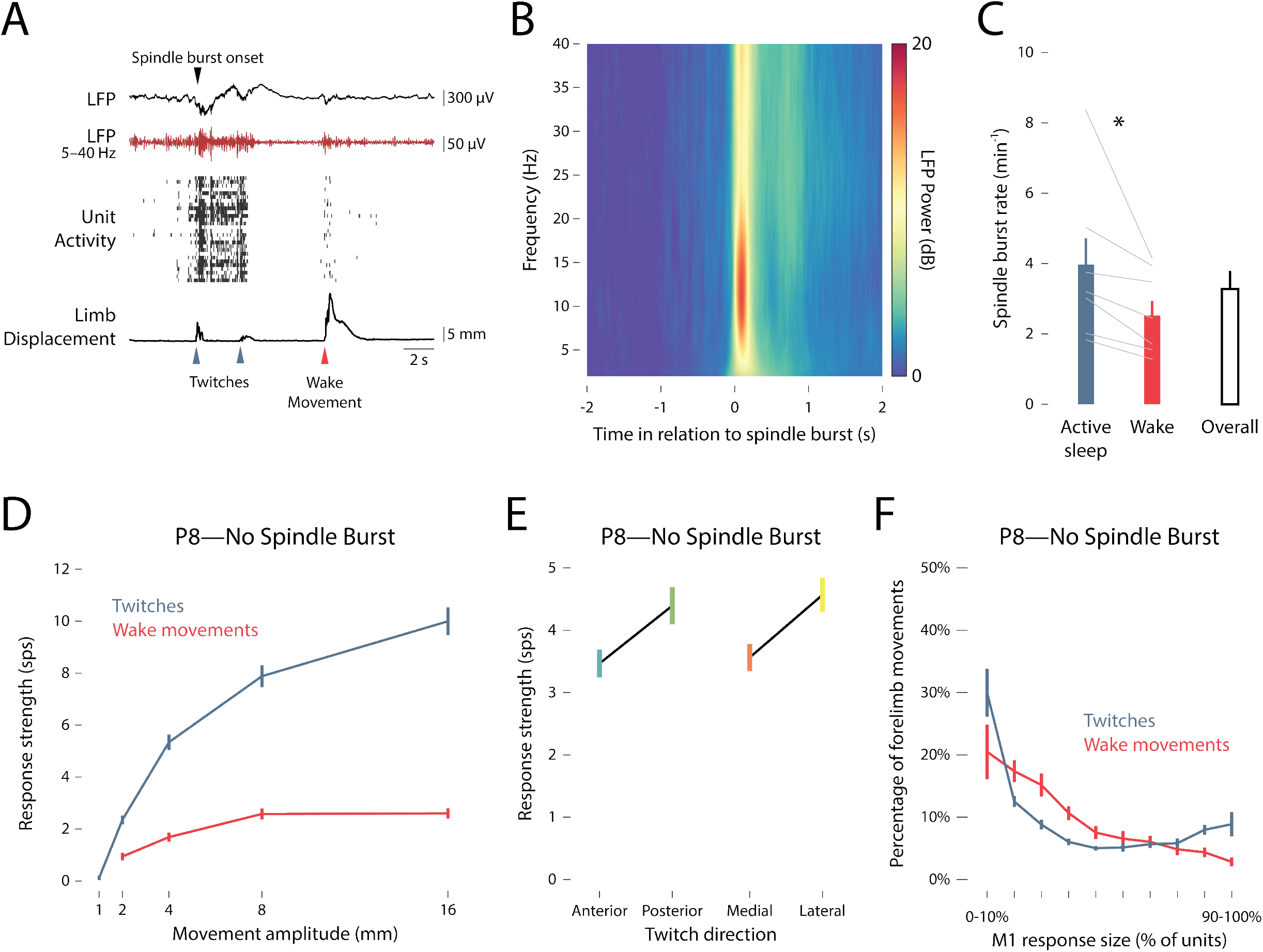
Kinematic tuning is not mediated by spindle bursts. (**A**) Representative data from a P8 rat. From top to bottom: A trace of the local field potential (LFP) in M1, including the onset of a spindle burst denoted with a black arrow; a filtered (5–40 Hz) LFP trace (red); unit activity in M1, with each row denoting a different unit and each vertical tick denoting an action potential; trace of forelimb displacement (black), with twitches and wake movements denoted by blue and red arrows, respectively. (**B**) Time-frequency spectrogram of LFP activity in M1, averaged across all P8 rats. The analysis was triggered on the onset of detected spindle bursts. (**C**) The rate of spindle bursts during active sleep (blue bar) and wake (red bar), as well as the overall rate (white bar) is shown for P8 rats. Gray lines show the values for individual pups. Asterisk denotes significance difference between sleep and wake (p < .05). (**D**) Same as in Fig 3B, but triggered only on movements that did not occur within ±0.5 s of a spindle burst. (**E**) Same as in Fig 4C, but triggered only on twitches that did not occur within ±0.5 s of a spindle burst. (**F**) Same as in Fig 6B, but triggered only on movements that did not occur within ±0.5 s of a spindle burst.

We determined the response strength within individual M1 units triggered on forelimb movements that did *not* occur within ±0.5 s of a spindle burst as a function of movement amplitude (Fig 7D) and movement direction (Fig 7E); response strength was largely unchanged compared with the earlier analyses for movement amplitude (Fig 3B) and movement direction (Fig 4C). M1 activity *across* units (population response) was similarly unaffected when triggered only on forelimb movements that did not occur within ±0.5 s of a spindle burst (Fig 7F). All together, these findings indicate that the kinematic properties of M1 units observed at P8, and the shift in kinematic properties at P12, are not mediated by spindle bursts.

## Discussion

Twitches are a distinct class of movement: They are brief and discrete, occur against a background of muscle atonia, and are highly diverse in their trajectories (i.e., amplitude and direction). As such, reafference arising from twitches is well-suited to drive somatotopically precise activity in developing sensorimotor cortex. By contrast, wake movements are typically prolonged and complex, involving sustained activity within and across multiple limb muscles; consequently, reafference arising from wake movements may be counterproductive to the development of precise forelimb somatotopy in sensorimotor cortex. These features of wake movements may help to explain why wake-related reafference is initially gated by the external cuneate nucleus (ECN; [4,6]), a primary recipient of ascending proprioceptive feedback from the forelimbs [24].

Consistent with previous studies [25-27], we found that active sleep is the predominant behavioral state in P8 rats (see S2 Fig). We also found at this age that twitches occurred three times more frequently than did wake movements and that twitches drove significantly more M1 activity than did wake movements. The increased frequency of twitches and their enhanced ability to drive M1 activity suggests that they play an important role in the development of M1’s somatotopic map. In turn, given that the somatotopic map at P8 and P12 exhibits the same topography as the motor map that will begin to emerge two weeks later [7], we propose that the somatotopic map serves as a reference that will guide motor outflow to the correct efferent target.

### Kinematic tuning in developing M1

We have demonstrated here in P8 rats that the processing of reafference in M1 is sensitive to the kinematic features of twitches—especially twitch amplitude. This is the second such demonstration of rate-coding of movement amplitude in developing rodent cortex: In P5 rats, we similarly found that larger whisker twitches more strongly drive neural activity in S1 barrel cortex [14]. Although spontaneous activity also occurs in the developing retina [10,20] and cochlea [28], it is not known whether this activity produces rate-coded responses in visual and auditory cortex, respectively.

It is not clear whether M1 amplitude-coding indicates tuning to the forelimb’s position in space *per se*, or simply tuning to the muscles that produce forelimb movements. For example, it is possible that larger forelimb movements tend to be produced by larger forelimb muscles (such as those at the shoulder), consequently triggering stronger reafferent activation of M1. If this were the case, one might expect that individual units would respond selectively to twitches of either large or small muscles (and thus either large or small movements). On the contrary, we found at P8 that individual units were responsive across a *range* of movement amplitudes. This observation suggests that each unit is not tuned to a specific muscle, but rather to movement amplitude across the entire limb.

Interestingly, once reafferent activity is conveyed to M1, it is modulated in a state-dependent manner that, again, resembles the state-dependent modulation found in S1 barrel cortex [14]. In fact, even though equally sized forelimb twitches and wake movements are presumably similar in terms of their patterns of forelimb muscle activation, twitches nonetheless produce substantially greater M1 activation (Fig 3B). This state-dependent amplitude tuning was particularly clear at the level of individual units: Of the 217 units recorded from at P8, 91% were tuned to twitch amplitude and 62% were tuned *exclusively* to twitch amplitude (data not shown).

At P8, nearly all M1 units were sensitive to twitch direction (see Fig 4). Assuming that twitches in different directions are caused by recruitment of different combinations of muscles, this finding suggests that M1 neurons at P8 are sensitive to the combination of forelimb muscles producing each twitch. However, EMG recordings of each forelimb muscle would be necessary to accurately assess the contributions of each muscle to M1 activity at this age. Notably, twitches with *uncommon* characteristics (e.g., large twitches, posterior/lateral twitches) produced the most unit activity, suggesting that neural activity in M1 is biased toward the detection of these twitches at P8. Thus, it may be that self-generated movements with relatively rare kinematic properties are amplified so as to ensure their continued representation in the forelimb region of M1. This issue could be explored further by monitoring the development of M1 tuning parameters in response to experimental manipulation of twitch amplitude or direction.

We also found that the informational content provided to M1 by twitch reafference increased from P8 to P12. At P8, amplitude and direction tuning were uniformly distributed across M1 units: 91% of units were tuned to twitch amplitude (Fig 3E) and 70–73% of units were tuned to a single twitch direction (Fig 4E). Thus, when considering all units together, the kinematic information about twitches was represented redundantly within M1. Redundancy decreased by P12: The number of amplitude-tuned units decreased to just 17% of all units, and at most 49% of units were tuned to a single direction. This decrease in redundancy, along with the increased responses of M1 units to small twitches at P12 (S3B), indicates an increase in informational content provided by twitches at P12. Therefore, whereas the system at P8 seems to prioritize the detection of self-generated movements, the system at P12 seems to prioritize the most efficient and informative representation of those movements. Accordingly, P12 may represent the beginning of a new developmental phase in which reafferent activity in M1 becomes organized within more complex, sparse networks.

Finally, spindle bursts, a predominant thalamocortical oscillation in neonatal rats [14,20-22], were too infrequent at P8 to mediate the observed effects of movement amplitude and direction on M1 activity (Fig 7C). Only 10% of forelimb movements coincided with spindle bursts, and exclusion of those movements from analysis did not alter the M1 response profiles (Fig 7D–F). Although one proposed role for spindle bursts is to strengthen developing thalamocortical circuitry [21,22,29], it is unclear how spindle bursts, compared with spiking activity alone, are contributing to this process at P8. However, because the rate of spindle bursts peaks during the first postnatal week and decreases thereafter [20,23,30], P8 may be beyond the age for properly identifying the functional contributions of spindle bursts to M1 development.

### Population activity in developing M1

At P8, population-level activity in M1 is discontinuous, occurring primarily in discrete, correlated bursts of unit activity separated by periods of silence (Fig 5A). Neural activity in S1 [31,32] and primary visual cortex (V1; [33]) is similarly discontinuous at this age. It has been hypothesized that discontinuous activity helps to maximize the detection of spontaneous peripheral activity (see [34]), thereby aiding in the activity-dependent development of sensory networks [10-12].

At P12, however, correlated activity disappears as background activity, sparsity (a measure of the uniform distribution of action potentials), and entropy (a measure of informational capacity) increase. Such *sparsification* of cortical activity in M1 also occurs around P12 in S1 [31,32] and V1 [33]. Sparsification of cortical activity is timed contemporaneously with the sudden emergence of local cortical inhibition [4,35,36]. Moreover, recent modeling studies suggest that developmental changes to the balance of inhibitory and excitatory processes in cortex contribute to the sparsification of cortical activity at P12 [37,38]. Finally, given that inhibitory interneurons strongly modulate cortical sensory processing in adulthood [39,40], the development of inhibition in M1 may explain the developmental reduction in redundant tuning properties observed here at P12.

It is possible that the onset of sparsification in M1 depends on the prior development of M1’s forelimb representation via sensory experience (i.e., reafference). Consequently, perturbing early sensory experience may disrupt the onset of sparsification in M1. Indeed, in neonatal mice, dark rearing (an example of sensory deprivation) delays sparsification in V1 by 2–3 days [33]; in contrast, whisker plucking in neonatal mice does not delay sparsification in S1 barrel cortex [31]. Given such conflicting results, more work is needed to clarify the conditions under which cortical sparsification is affected by early sensory experience.

Surprisingly, the distribution of M1’s population response to forelimb movements shifted from a roughly exponential to a roughly normal distribution over just four days (Fig 6B). This shift in response distributions meant that M1 at P12 was less likely to exhibit an all-or-none response to a forelimb movement and more likely to exhibit a response comprising approximately half of the M1 units. This absence of redundant population responses at P12 implies that forelimb movements provide more informational content to M1 at this age. This new phase in M1 development in which reafferent input is more reliably detected and more informationally dense may serve to improve the effectiveness of M1’s sensory map as a reference for its later-developing motor map.

The increase in variability in kinematic coding and robust changes to population-level activity shown here foreshadows M1 functioning in adulthood: Adult M1 neurons are exquisitely versatile and heterogenous, with each neuron able to simultaneously represent multiple kinematic (as well as temporal) features of movement in both Cartesian and body-centered reference frames [41-43]. M1’s complexity in adulthood is partially supported by “sparse coding”—in which complex sensory input is reflected in precise, energetically efficient responses [44]. The transition to more complex sensory coding at P12, then, signifies an important transition toward the development of complex network properties, such as sparse coding, that will eventually enable M1 to produce complex movements and participate in motor learning [17,18].

## Methods

### Resource Availability

#### Lead Contact

Further information and requests for resources and code should be directed to, and will be fulfilled by, the lead contact, Dr. Mark Blumberg.

#### Materials Availability

This study did not generate new unique reagents.

#### Data and Code Availability

Raw data (action potential timestamps, behavioral event timestamps, and forelimb position time-series) will be made available upon request. Select custom MATLAB scripts used here can be found on Github (https:\\www.github.com\XXXXX). Additional scripts and data used will be made available upon request.

### Experimental Model and Subject Details

Sprague-Dawley Norway rats (*Rattus norvegicus*) were used at P8–9 (n = 8, body weight: 18.8 ± 0.4 g; hereafter referred to as P8) and P12–13 (n = 8, body weight: 30.6 ± 0.7 g; hereafter referred to as P12). Equal numbers of males and females were used and all subjects were selected from separate litters. In total, 37 single-units and 180 multi-units were isolated at P8 and 106 single-units and 144 multi-units were isolated at P12. In preliminary analyses, multi-units showed parsimonious results with single units and thus were included in all analyses (217 and 250 combined units at P8 and P12, respectively).

Animals were housed in standard laboratory cages (48 × 20 × 26 cm) on a 12:12 light-dark cycle, with food and water available ad libitum. The day of birth was considered P0 and litters were culled to eight pups by P3. All experiments were conducted in accordance with the National Institutes of Health (NIH) Guide for the Care and Use of Laboratory Animals (NIH Publication No. 80–23) and were approved by the Institutional Animal Care and Use Committee of the University of Iowa.

### Method Details

#### Surgery

As described previously [4,45], a pup with a visible milk band was removed from the litter and anesthetized with isoflurane gas (3–5%; Phoenix Pharmaceuticals, Burlingame, CA). A custom-made bipolar hook electrode (0.002-inch diameter, epoxy coated; California Fine Wire, Grover Beach, CA) was inserted into the nuchal muscle for state determination. Carprofen (5 mg/kg SC; Putney, Portland, ME) was administered as an anti-inflammatory analgesic. After removing the skin above the skull, an analgesic was applied topically (bupivacaine; Pfizer, New York, NY). The skull was dried with bleach. Vetbond (3M, Minneapolis, MN) was applied to the skin around the skull and a head-plate (Neurotar, Helsinki, Finland) was secured to the skull using cyanoacrylate adhesive.

A trephine drill bit (1.8 mm; Fine Science Tools, Foster City, CA) was used to drill a hole into the skull above the left forelimb representation of M1 (1.0 mm anterior to bregma, 2.2–2.5 mm lateral from the sagittal suture). Two smaller holes were drilled distally to the recording site for insertion of a thermocouple and reference/ground electrode. A small amount of peanut oil was applied over the recording site to prevent drying of the cortical tissue. Surgical procedures lasted approximately 15 min.

The pup was then secured to a custom-made head-restraint apparatus inside a Faraday cage, with the animal’s torso supported on a narrow platform. Brain temperature was monitored using a fine-wire thermocouple (Omega Engineering, Stamford, CT) distal to the M1 recording site. The pup was allowed to recover from anesthesia and acclimate for at least 1 h. The 1-h recording period did not begin until brain temperature was 36–37° C and the pup was cycling regularly between sleep and wake.

#### Electrophysiological Recordings

The nuchal EMG electrode was connected to a Lab Rat LR-10 acquisition system (Tucker Davis Technologies, Gainesville, FL). The EMG signal was sampled at approximately 1.5 kHz and high-pass filtered at 300 Hz.

A 16-channel silicon depth electrode (Model A4×4-3mm-100-125-177-A16; NeuroNexus, Ann Arbor, MI) was coated in fluorescent Dil (Vybrant Dil Cell-Labeling Solution; Life Techologies, Grand Island, NY) before insertion. The electrode was inserted 600–1000 µm into the forelimb representation of M1, angled 6° medially. A chlorinated Ag/Ag-Cl wire (0.25 mm diameter; Medwire, Mt. Vernon, NY) was inserted distal to the M1 recording site, serving as both a reference and a ground. The neural signal was sampled at approximately 25 kHz, with a high-pass (0.1 Hz) and a harmonic notch (60, 120, and 180 Hz) filter applied.

Electrode placement in the forelimb region of M1 was confirmed by manually stimulating the forelimb and observing exafferent neural activity. (Because forelimb stimulation also triggers activity in primary somatosensory cortex, histology was performed to further confirm electrode placement in M1; see below.) Neural activity from M1 was recorded for 1 h using SynapseLite (Tucker Davis Technologies) while the animal cycled between sleep and wake states.

#### Video Collection

In order to digitally reconstruct forelimb movements in three dimensions, video of the forelimb was recorded from front and side camera angles using two Blackfly-S cameras (FLIR Integrated Systems; Wilsonville, Oregon). Video was collected in SpinView (FLIR Integrated Systems) at 100 frames/s, with a 3000-μs exposure time and 720×540 pixel resolution. The two cameras were hardwired to acquire frames synchronously and were initiated using a common software trigger.

The synchronization of video and electrophysiological data was ensured by using an external time-locking stimulus. A red LED controlled by SynapseLite (Tucker Davis Technologies) was set to pulse every 3 s for a duration of 20 ms. The LED was positioned to be in view of both cameras. Each video was analyzed frame-by-frame with custom Matlab scripts to ensure an equal number of frames between LED pulses. Although infrequent, when the number of frames between pulses was less than expected, the video was adjusted by duplicating and inserting one adjacent frame at that timepoint so as to preserve timing across the recording. Thus, all videos were ensured to be time-locked to the electrophysiological data within 10 ms.

#### Histology

At the end of the recording period, the pup was euthanized with ketamine/xylazine (10:1; >0.08 mg/kg) and perfused with 0.1 M phosphate-buffered saline, followed by 4% paraformaldehyde. The brain was extracted and post-fixed in 4% paraformaldehyde for at least 24 h and was transferred to a 30% sucrose solution 24–48 h prior to sectioning.

In order to confirm the electrode’s location within the forelimb representation of M1, the left cortical hemisphere was dissected from the subcortical tissue and flattened between two glass slides (separated using two pennies) for 5–10 min. Small weights (10 g) applied light pressure to the top glass slide. The flattened cortex was sectioned tangentially to the pial surface. A freezing microtome (Leica Microsystems, Buffalo Grove, IL) was used to section the cortex (80-µm sections). Free-floating sections were imaged at 2.5x using a fluorescent microscope and digital camera (Leica Microsystems) to mark the location of the DiI.

Electrode placement in the forelimb region of M1 was confirmed by staining cortical sections for cytochrome oxidase (CO), which reliably delineates the divisions of primary sensory cortex at this age [46]. The M1 forelimb representation is immediately medial to (and partially overlaps) the primary sensory forelimb representation (see Fig 1B–C). Cytochrome C (0.3 mg/mL; Sigma-Aldrich), catalase (0.2 mg/mL; Sigma-Aldrich), and 3,3’-diaminobenzidine tetrahydrochloride (DAB; 0.5 mg/mL; Spectrum, Henderson, NV) were dissolved in a 1:1 dilution of 0.1 M phosphate buffer and distilled water. Sections were developed in 24-well plates on a shaker (35–40°C, 100 rpm) for 3–6 h, rinsed in phosphate-buffered saline, and mounted on a glass slide. The stained sections were imaged at 2.5x or 5x magnification and composited with the fluorescent image to confirm the electrode tract within the forelimb region of M1.

#### Behavioral State and Forelimb Movements

As described previously [3,4], nuchal EMG and behavior were used to assess behavioral state (the experimenter was blind to the neurophysiological record while scoring behavior). The wake state was defined by the presence of high-amplitude movements of the limbs against a background of high nuchal muscle tone. Active sleep was defined by the presence of discrete myoclonic twitches of the face, limbs, and tail against a background of nuchal muscle atonia.

Forelimb movements were quantified using DeepLabCut (DLC), a markerless tracking solution that uses a convolutional neural network to track features (e.g., limbs) of animals in a laboratory setting [15,16]. At least 200 manually labeled frames (tracking the wrist of the right forelimb) were used to initially train the network. After the initial training, newly analyzed frames with marker estimates that were deemed inaccurate were re-labeled and used to re-train the neural network until satisfactory tracking was achieved. Separate neural networks were trained for the front-facing and side-facing camera angles. The networks reached a training root mean square error (RMSE) of 0.18 and 0.19 mm and a test RMSE of 0.28 and 0.42 mm, respectively.

Forelimb twitches and wake movements were identified using custom Matlab scripts. Although infrequent (0.5 ± 0.1% of frames), individual frames in which the wrist position was associated with a low confidence value (<.80, identified by DLC) were removed and linearly interpolated from the position data of adjacent frames. Forelimb position data was derived to obtain forelimb velocity, and forelimb twitches and wake movements were detected by peak detection of forelimb velocities reaching 2x the standard deviation of quiet periods for twitches, and 3x the standard deviation of the quiet period for wake movements. All movements were required to be preceded by a 250-ms period of quiescence in the forelimb. Every forelimb twitch and wake movement was visually confirmed by the experimenter to identify and discard false positives.

For the analysis of movement amplitude, the forelimb position was summed across the x-, y-, and z-dimensions using the Pythagorean theorem (Fig 2A). The peak amplitude was measured as the difference between the point of maximum displacement (using a shared time point for each dimension) and the median baseline position from -0.5 to 0 s before the initiation of a twitch or wake movement. Because twitch amplitude follows an exponential distribution (see Fig 2B), movements were sorted into logarithmic bins of 0–1, 1–2, 2–4, 4–8, and 8–16 mm.

For the analysis of movement direction, the forelimb position was transformed into polar coordinates along the rostral-caudal and medial-lateral axes (see Fig 2C). Movements were sorted into anterior-posterior or medial-lateral bins. Because wake movements occur during sustained bouts of continuous activity and regularly involve multi-directional trajectories, they could not be analyzed for direction-tuning in M1.

#### Spindle Bursts

As described previously [14], the neural signal was band-pass filtered at 5–40 Hz with a stopband attenuation of -60 dB and a 1-Hz transition gap. A Hilbert transformation was applied to the filtered waveform, and spindle burst onset was defined as the first point at which the waveform amplitude exceeded the median plus two standard deviations. Spindle bursts were defined as having a minimum duration of 150 ms.

#### Spike Sorting

SynapseLite files were converted to binary files using custom Matlab scripts and sorted with Kilosort [47]. Briefly, the 16 channels of neural data were whitened (covariance-standardized) and band-pass filtered (300–5000 Hz) before spike detection. Next, template matching was implemented to sort the event waveforms into clusters. The first-pass spike detection threshold was set to 6 standard deviations below the mean and the second-pass threshold was set to 5 standard deviations below the mean. The minimum allowable firing rate was set to 0.01 sps and the bin size for template detection set to 656,000 sample points for P8 animals and 262,400 sample points for P12 animals (approximately 27 s and 11 s, respectively). All other Kilosort parameters were left at their default values.

Clusters were visualized and sorted in Phy2 [48]. Putative single units (elsewhere referred to as “single-units”) were defined as having (1) spike waveforms that reliably fit within a well-isolated waveform template, (2) a distribution along seven principal components that was absent of clear outliers, and (3) an auto-correlogram with a decreased firing rate at time-lag 0 (indicative of a single-unit’s refractory period).

Clusters meeting the first two criteria but not the third were considered multi-units. Any cluster with a waveform template indicative of electrical noise, a significantly low firing rate (< 0.01 sps), or amplitude drift across the recording period was discarded.

#### Data Shuffling

Two shuffling procedures were used to approximate the null hypotheses in (1) kinematic analyses and (2) population-activity analyses. First, for all kinematic analyses (i.e., those performed in Fig 3, 4, and 6C–D), twitches and wake movements were randomly selected in proportion to the number of movements used in the main analysis. 100 analyses were performed using randomly selected forelimb movements and the 99% confidence interval was computed to approximate the distribution of the null hypothesis. For example, in Fig 3B, twitches were randomly selected (i.e., amplitude was ignored, but direction was controlled for) and analyzed 100 times and the 99% confidence interval was taken to obtain the shuffled data.

Second, for all population-activity analyses (i.e., those performed in Fig 5 and 6B), the inter-spike intervals of each spike train were resampled and used to create shuffled spike trains. This procedure conserves each unit’s overall firing rate and temporal dynamics [49,50]. Analyses using the resampled spike trains were repeated 100 times and the 99% confidence interval computed to obtain the shuffled data.

#### Population Analyses

The onset of movement periods was defined as the onset of twitches and wake movements described above. The offset of each movement period was defined as the time-point 250 ms after the displacement of the forelimb returned below the threshold for detection (described above) for the first time. Therefore, each movement period was followed by 250 ms to account for reafference arising from each movement. The onset of non-movement periods was defined as the offset of movement periods, and therefore were preceded by 250 ms of quiescence in the limb. The offset of the non-movement periods was defined as the initiation of the next movement. All non-movement periods less than 250 ms and all periods that overlapped were discarded from analysis (<2% of the duration of any recording).

Sparsity was defined according to [51]

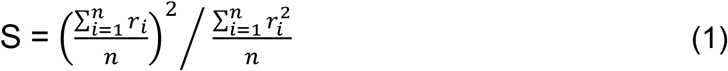

where **n** represents the number of forelimb movements of a given pup and ***r***_***i***_ represents the firing rate of a given unit from 0–250 ms after the ***i***th forelimb movement.

Entropy (of a discrete random variable) was defined according to [52]

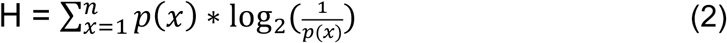

where **n** represents the number of possible states for a unit (based on its firing rate) and ***p***(***x***) represents the probability distribution of a given unit in that state. Firing-rate data were discretized according to procedures outlined in [53]. Briefly, a unit’s firing rate was calculated in 250-ms time windows, and each window was assigned to one of three possible states based on a uniform-width binning of the firing rate distribution.

For correlation analyses, movement and non-movement periods were broken down into 250-ms time windows and a Pearson correlation coefficient of the firing rate was computed across time windows for every possible unit-unit pair. All Pearson coefficients for an animal were averaged together before comparison between P8 and P12.

*Population response* was defined as the percentage of all units of a given pup that were “responsive” to a given forelimb movement. A unit was considered responsive if its firing rate significantly increased (i.e., above the 95% confidence interval of baseline firing rate) after a forelimb movement; otherwise, a unit was considered unresponsive.

### Quantification and Statistical Analysis

#### Statistical Analysis

All data were tested for normality using the Shapiro-Wilk test, for equal variance using Levene’s test (for between-subjects variables), and for sphericity using Mauchly’s test (for within-subjects variables with >2 groups) prior to analysis. In analyses in which the variance between groups was not equal, a pooled error term was not used when generating simple main effects and post-hoc tests. In analyses in which sphericity was violated, a Huynh-Feldt correction was applied to the degrees of freedom. Probabilities and r^2^ values were arc-sin transformed prior to analysis. The mean and standard error of the mean (SEM) are used throughout as measures of central tendency and dispersion, respectively.

All analyses were performed as independent t tests, or two- or three-way mixed design ANOVAs. Throughout the Results section, simple main effects were only reported if the interaction term was significant. All main effects, interactions, simple main effects, and the results of any additional statistical tests can be found in S4 Table.

In all two- and three-way ANOVAs, an adjusted partial eta-squared was used as an estimate of effect size that corrects for positive bias due to sampling variability [54]. For all t tests and one-way ANOVAs, an adjusted eta-squared estimate of effect size was reported.

## Acknowledgments

This research was supported by a grant from the National Institute of Child Health and Human Development (R37-HD081168) to M.S.B.

## Author Contributions

Conceptualization, R.M.G., J.C.D, G.S., and M.S.B.; Methodology, R.M.G., J.C.D, G.S., and M.S.B.; Software, R.M.G. and J.C.D.; Validation, R.M.G.; Formal Analysis, R.M.G.; Investigation, R.M.G.; Resources, G.S. and M.S.B.; Data Curation, R.M.G.; Writing - Original Draft, R.M.G.; Writing—Reviewing and Editing, R.M.G., J.C.D., G.S., and M.S.B.; Visualization, R.M.G. and M.S.B.; Supervision, G.S. and M.S.B.; Project Administration, G.S. and M.S.B.; Funding Acquisition, G.S. and M.S.B.

---

**S1 Fig.**
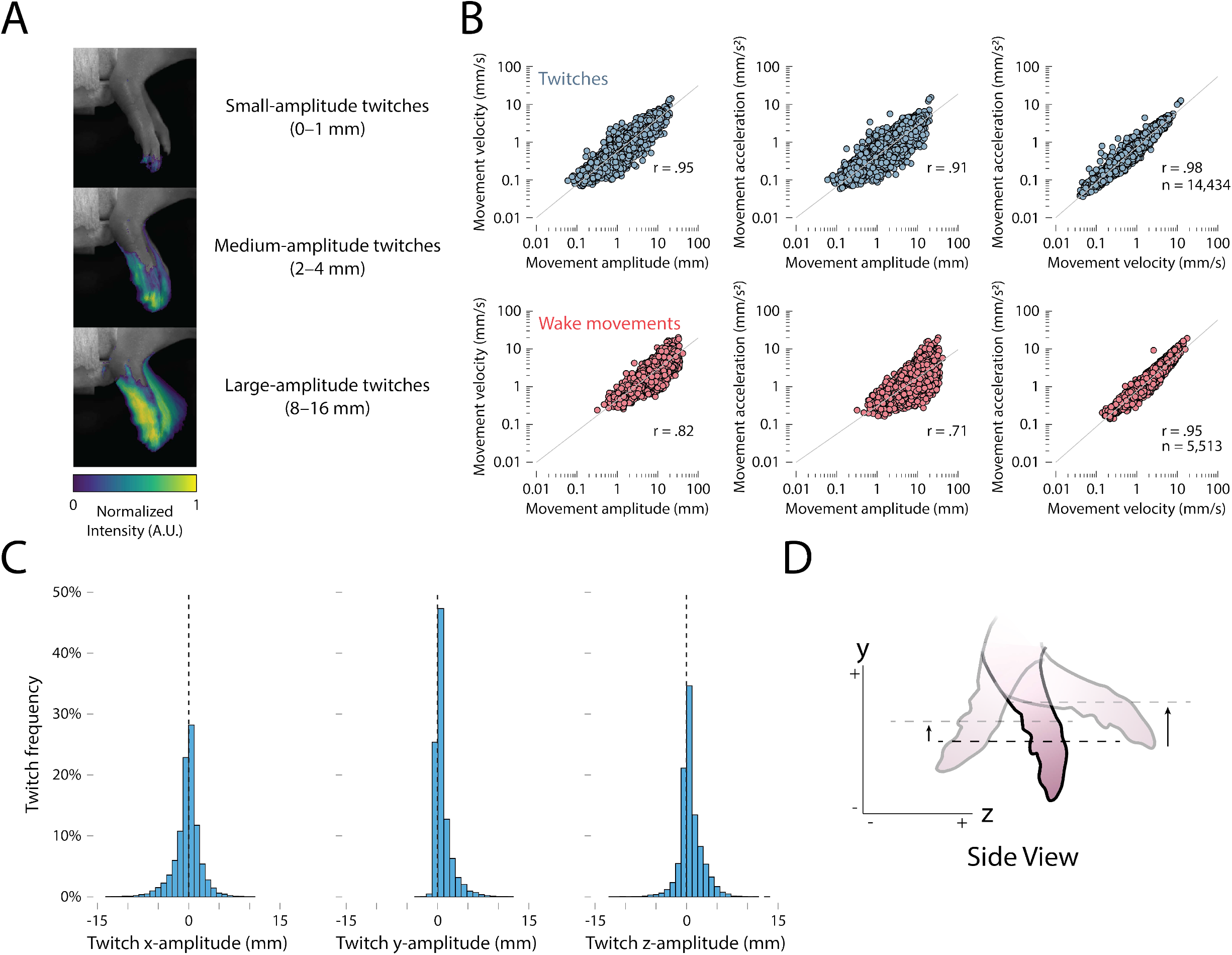
Additional kinematic data for forelimb movements. (**A**) The average change in pixel intensity within the region-of-interest across 50 small-amplitude (top), medium-amplitude (middle), and large-amplitude (bottom) twitches is shown as a heatmap. Smaller twitches primarily reflect displacement of the digits and larger twitches typically reflect displacement of the digits, wrist, and elbow. (**B**) Bivariate correlations for all twitches (top row) and wake movements (bottom row) for movement velocity vs. movement amplitude (left column), movement acceleration vs. movement amplitude (center column), and movement acceleration vs. movement velocity (right column). In each plot, movements are pooled across age. All r values are significant at p < .05. (**C**) Relative frequency histograms depict the displacement of twitches (data from P8 and P12 rats combined) along the x- (left), y- (center), and z- (right) dimensions. Note the asymmetrical distribution of twitches in the positive y-dimension, which prompted the exclusion of this dimension from further direction analyses. (**D**) Side view of right-forelimb movements to show how pendular motion produces only a positive y-displacement.

**S2 Fig.**
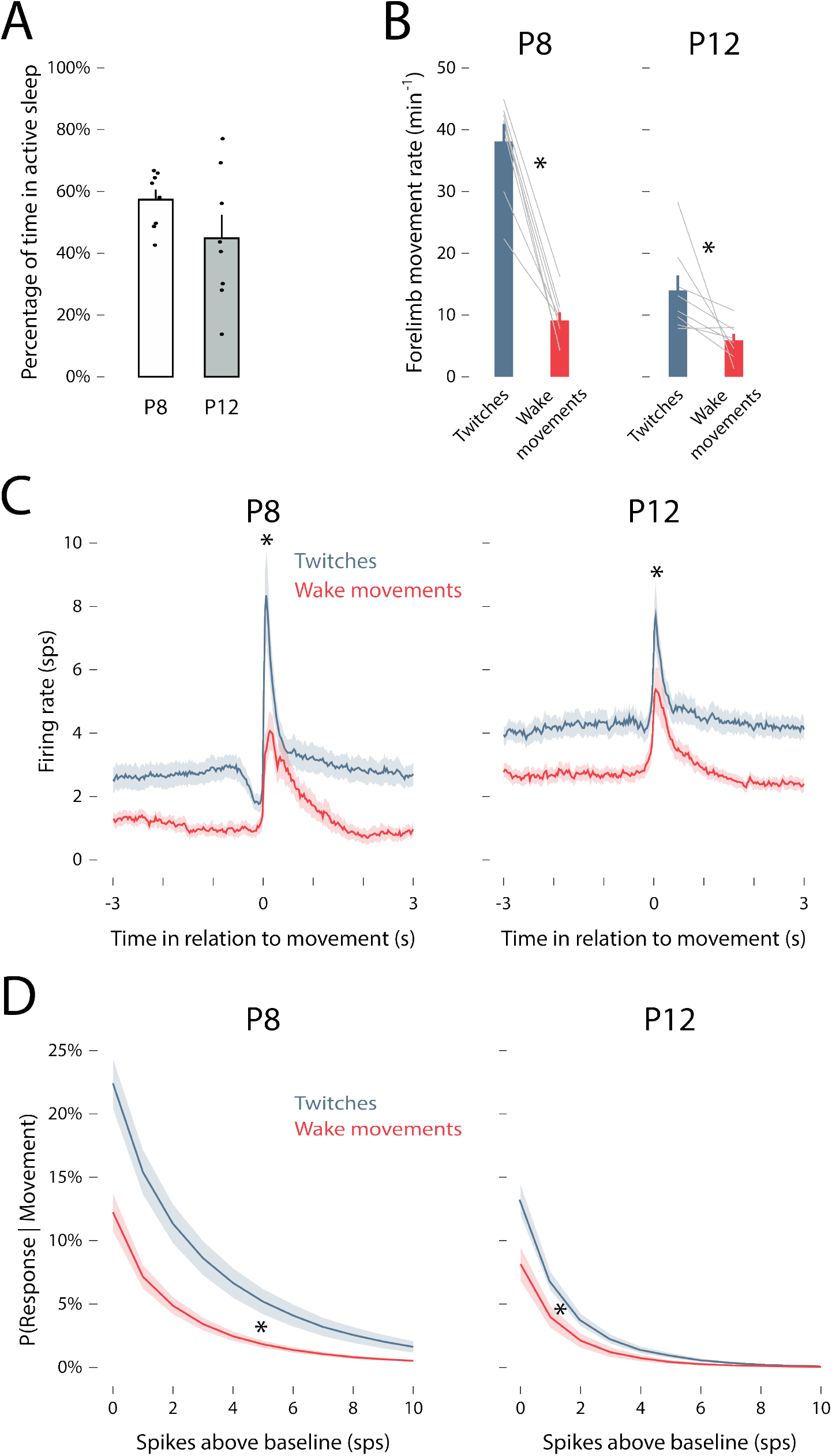
Twitch and wake movements trigger reafferent responses in M1. (**A**) Mean (±SEM) percentage of time spent in active sleep at P8 (white) and P12 (gray). Black dots show values for individual pups. (**B**) Mean (±SEM) rate of forelimb movements for twitches during active sleep (blue) and movements during wake (red) at P8 (left) and P12 (right). Gray lines show values for individual pups. Asterisks denote significant difference between twitches and wake movements (p < .05). (**C**) Peristimulus time histograms of mean firing rate of M1 units in relation to twitches (blue) and wake movements (red) at P8 (left) and P12 (right). Asterisks denote significant difference between responses to twitches and wake movements (p < .05). (**D**) Mean (±SEM) probability of an M1 response (individual unit) given a twitch (blue) or wake movement (red) as a function of spiking threshold (i.e., firing rate above that unit’s baseline firing rate). Higher spiking thresholds yield smaller response probabilities. Asterisks denote significant difference between twitches and wake movements (p < .05).

**S3 Fig.**
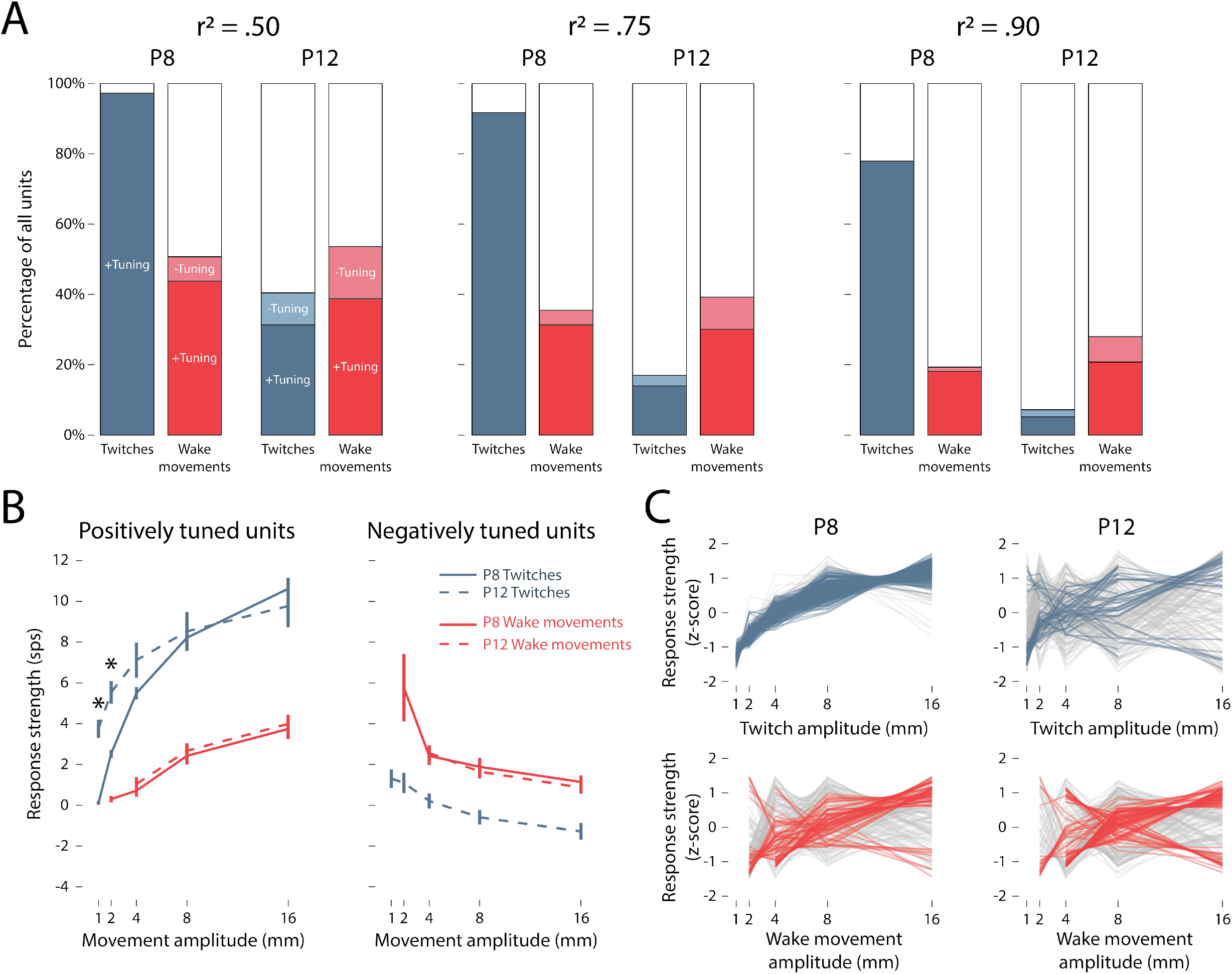
Additional aspects of the relationship between movement amplitude and M1 unit responses. (**A**) As in Fig 3E, the percentage of units that are positively tuned to movement amplitude for twitches (dark blue) and wake movements (dark red), negatively tuned for twitches (light blue) and wake movements (light red), or not tuned (white). From left to right, two r2 thresholds are shown below (.50) and above (.90) the threshold of .75 used in Fig 3E. (**B**) As in Fig 3B, mean (±SEM) response strength is shown as a function of movement amplitude. Solid and dashed blue lines denote P8 and P12 twitches, respectively; solid and dashed red lines denote P8 and P12 wake movements, respectively. Positively and negatively tuned units were identified using an r2 threshold of .75. Asterisks indicate significant difference between P8 and P12 (p < .05). (**C**) As in Fig 3B, change in response strength (Δfiring rate in relation to baseline, z-scored) is shown for individual M1 units in response to twitches (blue) and wake movements (red) at P8 and P12. Positively and negatively tuned units are indicated in blue (twitches) and red (wake movements); untuned units are indicated in gray.

**S4 Table.**
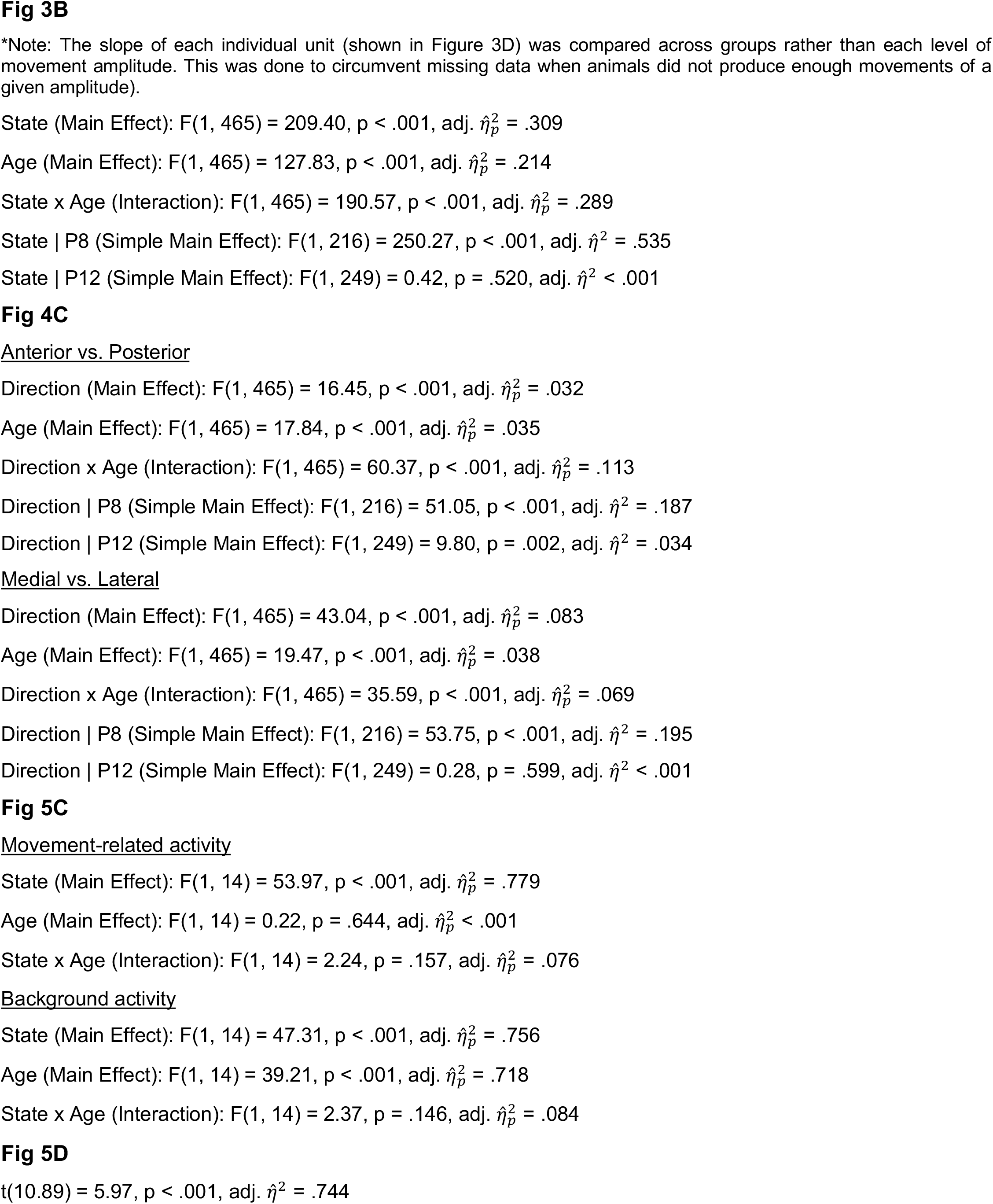
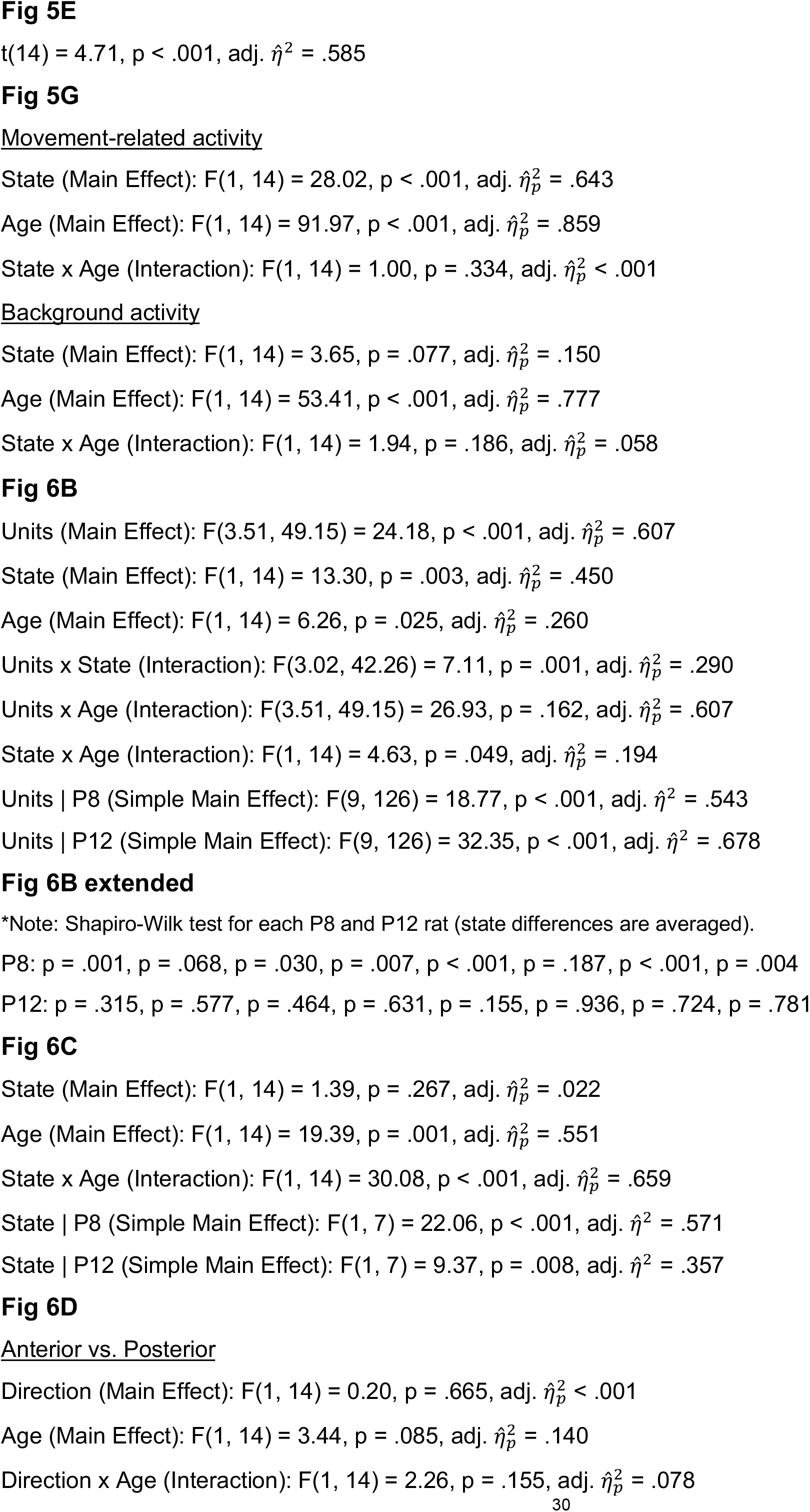
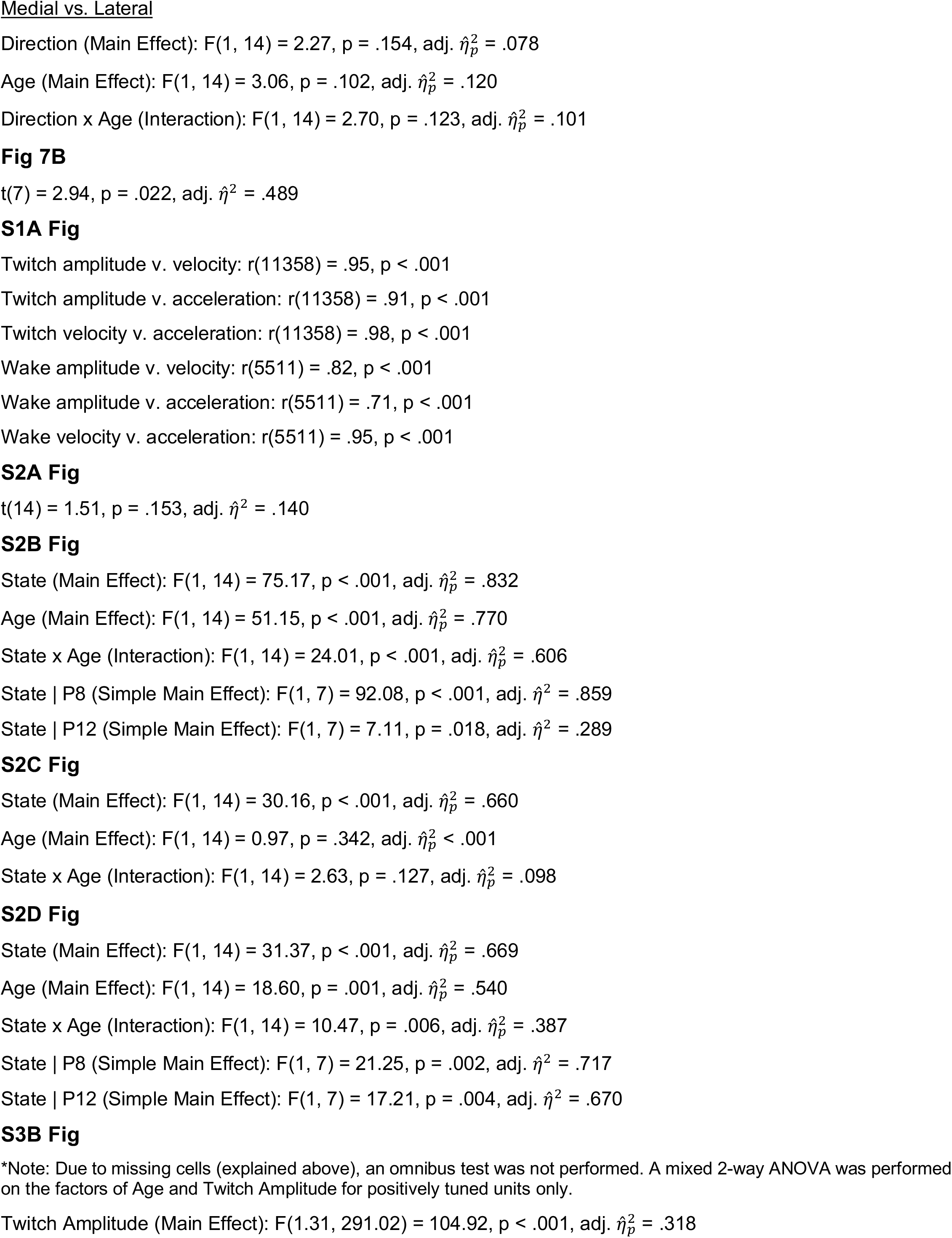
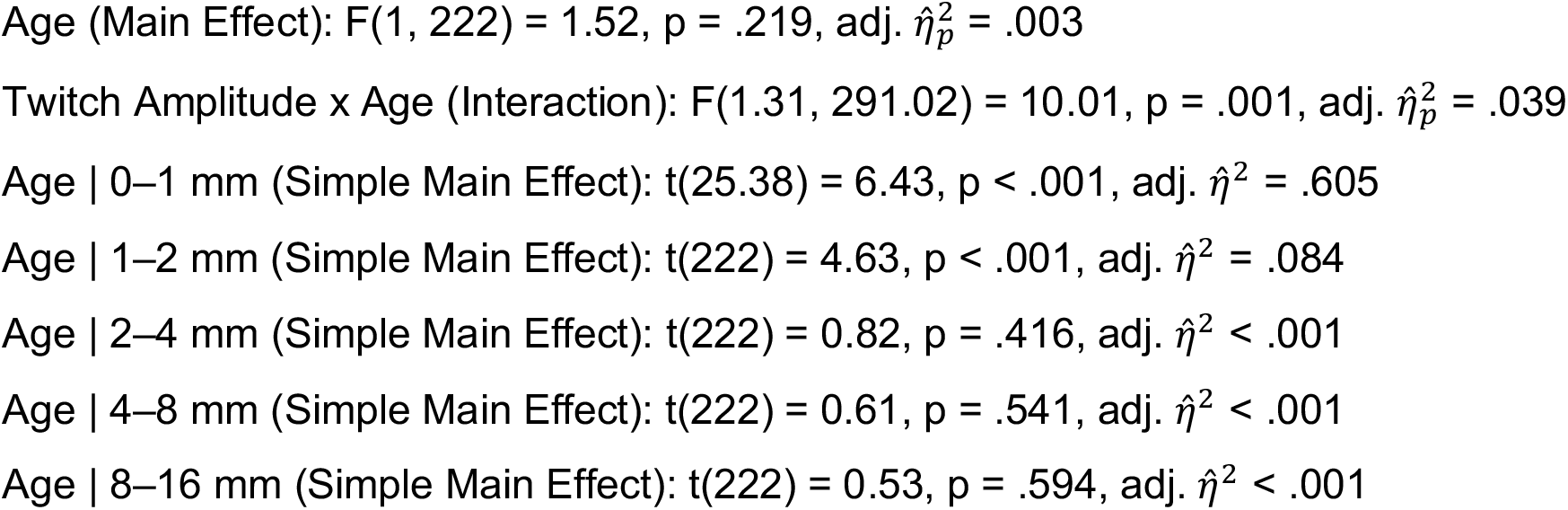

